# Short lifespan is one’s fate, long lifespan is one’s achievement: lessons from *Daphnia*

**DOI:** 10.1101/2024.03.30.587428

**Authors:** Thomas Beam, Mchale Bright, Amelia C. Pearson, Ishaan Dua, Meridith Smith, Ashit K. Dutta, Shymal C. Bhadra, Saad Salman, Caleb N. Strickler, Cora E. Anderson, Leon Peshkin, L. Y. Yampolsky

## Abstract

Studies of longevity and senescence rely on baseline life expectancy of reference genotypes measured in standardized conditions such as food level and group size. Variation in baseline lifespan data across labs and protocols and among genotypes can make longevity intervention studies difficult to compare, particularly when GxE interactions exist. Furthermore, extending the lifespan of a short-lived genotype or of any genotype under suboptimal conditions may be of a lesser theoretical and translational value than extending the maximal possible lifespan. *Daphnia* is rapidly becoming a model organism of choice for longevity research complementing data obtained on traditional models. In this study we report baseline longevity of several genotypes (parthenogenetic clones) of a long-lived species *D. magna* under a variety of laboratory protocols, aiming to document the highest possible lifespan, factors reducing it, and physiological parameters that change with age and correlate with longevity. Combining data from 25 different experiments across two labs we report strong differences among clones of different geographic origin, moderate effects of group size and medium composition on longevity, and strong GxE with respect to food level. Specifically, short-lived clones that tend to originate from small intermittent habitats show little or no caloric restriction (CR) longevity extension, while long-lived ones expand their lifespan even further when maintained at 25% of the *ad libitum* food. We find no evidence of any trade-offs between longevity and fecundity across clones or correlations with age-specific feeding rate. We find that in the short-lived, CR non-responsive clones show little correlation between longevity and two measures of lipid peroxidation (LPO: lipid hydroperoxides and MDA abundance). In contrast, the long-lived, CR-responsive clones show a positive longevity correlation with lipid hydroperoxide abundance at any age, and a negative correlation with MDA concentration measured at about median lifespan. This indicates differences among genotypes in longevity-related accumulation of LPO targets, efficiency of detoxification of LPO products, and/or their effects on longevity. Our observations support the hypothesis that a long lifespan can be affected by food availability and levels of oxidative damage, while genetically determined short lifespan remains short regardless. We suggest a set of condition and genotypes to be used as a reference for longevity studies in *Daphnia*.

## Introduction

The plankton microcrustacean *Daphnia* is a promising model organism for longevity and senescence studies (Dudycha et al., 2003; Constantinou et al. 2019). It reproduces by cyclic parthenogenesis, which creates a possibility to maintain genetically uniform and yet outbred clones, thus reducing within-cohort heterogeneity issues. Its semi-transparent body provides a possibility for in-vivo microscopy and fecundity measurements. Large body-to-water interface area and constant movement of water through the filtering apparatus and the guts make delivery of many potential longevity intervention drugs from the medium into tissues readily possible. Yet, so far *Daphnia* longevity studies have been suffering from the lack of a set standardized methods, in particularly in terms of isolates used, media composition, maintenance protocol and, importantly, food concentrations used.

Longevity studies, including longevity extension intervention experiments depend on accurate estimation and standardization of the background (control) lifespan (Spiridonova et al. 2024) and suffer from a significant variation in measured effects across treatments and labs, as wells as across species, genotypes, and sexes of the study organisms (Finch and Ruvkun 2001; Ziehm et al. 2013; Lucanic et al. 2017; Banse et al. 2019; Bartke et al. 2019; Yuan et al. 2020; Urban et al. 2021; Spiridonova et al. 2024). This virtually eliminates the possibility to compare life-extending effects of chemical or dietary interventions measured in different studies, as control (untreated) groups’ longevity can differ greater than anticipated intervention effects. In some high-profile cases what had been previously thought to be a well-supported result has been proven to be an artifact of lack of standardization of genetic background (Burnett et al. 2011). In the case of Drosophila data revisited by Burnett et al. (2011) the confounding factor making the originally used controls inappropriate were the effects of transgenic line generating procedure, but of course a similar problems can arise due to natural background variation in longevity-related traits.

Furthermore, arguably longevity intervention studies are the most useful, both from the theoretical and the potential translational standpoint, if they are conducted under conditions allowing highest possible lifespan in the control group (for both genetic and environmental reasons), and therefore any interventions are truly extending lifespan (with potential translational significance), and not compensating for genetic or environmental deficiencies. In fact, in case of genetic deficiencies in inbred genotypes, even that is unlikely, as each isolate is probably showing shortened lifespan due to being homozygous for a unique (set of) harmful recessive allele(s).

Genetic differences for lifespan are well documented in all organisms studied (Finch and Ruvkun 2001; Yuan et al. 2020; Kaya et al. 2021). They include naturally occurring variation (Kaya et al. 2021), including variation among nature-derived inbred lines (Mackay 2002; Dick et al. 2011; Mackay et al. 2012) and differences between evolved and control lineages (Rose 1984; Doroszuk et al. 2012). While informative for future intervention studies due to the possibility of elucidating mechanisms behind unusually long lifespan (Parker et al. 2020), genetic variants of lifespan also constitute a challenge when choosing the right isolate for intervention experiments (Lucanic et al. 2017). Different labs may work with different isolates resulting in different baseline longevity, making across-labs comparison difficult and, for the above mentioned reasons, positive results both unlikely and, if any, idiosyncratic to a particular genotype. If genotype-by-environment interactions for longevity are also present (Dick et al 2011; Huang et al. 2020; Rohde et al. 2021; Pallares et al. 2023; Simons and Dobson 2023), choosing different isolates as study systems may also result in little reproducibility of intervention results. *Daphnia* as a model organism is no exception, with a significant clonal variation in lifespan known between sister species (Dudycha et al. 2003), among geographically distinct populations (Dudycha and Hassel 2013; Lohr et al. 2014; Coggins et al. 2021; Ukhueduan et al. 2022) and within a population (Yampolsky and Galimov 2005). A significant portion of geographic lifespan variation is consistently associated with the difference between clones originating from small, temporary ponds and polls and those from larger, permanent ponds and lakes (Dudycha and Hassel 2013; Lohr et al. 2014). Thus, there is both an opportunity to further investigate the evolutionary and genomics causes of such variation and the need for standardization of choice of isolates for longevity intervention studies.

The same reasoning applies to environmental determinants of lifespan and their potential interactions with longevity interventions. While temperature, humidity, mineral composition of the environment, crowding, and social structure of cohorts may have strong effects on cohorts’ longevity in various organisms, perhaps the most pervasive environmental parameter affecting lifespan (and likely to interact with longevity interventions) is the diet. Caloric restriction (CR) effect on lifespan has been very well documented in a variety of organisms (Fontana et al. 2010; Fontana and Partridge, 2015), including yeasts (Wei et al. 2008), nematode worms (Kauffman et al. 2010; Loo et al. 2023), flies (Pletcher et al. 2005; Jin et al. 2020), rodents (Liao et al. 2010) and primates (Lane et al. 1997). *Daphnia* species were among the first organisms in which lifespan extension under caloric restriction has been discovered (Ingle 1933; Ingles et al. 1937) and it has been extensively documented since (Lynch and Ellis 1983; Lynch 1989; Pietrzak et al. 2010). However, a more complex pattern has emerged. First, it has been demonstrated that in an across-species comparison, a long-lived *D.pulicaria* shows much stronger life-extension CR effect than a short-lived sibling species *D.pulex* (Dudycha et al. 2003), although the effect is present in both. Further studies, however, either failed to replicate this observation (Latta et al. 2011) or failed to observe any CR effect at all (Schwartz et al. 2016). These findings raised the question whether *Daphnia* CR effect is universal or highly condition, species, or genotype-dependent. Genetic differences in the strength or even existence of the caloric restriction effect on longevity demonstrated, among other organisms, in *Drosophila* (Dick et al. 2011; Mackay et al. 2012; Zhu et al. 2014; Simons and Dobson 2023) and mice (Liao et al. 2013) have been of a considerable interest, as the varying strength of the CR effect on different genetic background may elucidate possible mechanisms of lifespan extension under CR conditions (Liao et al. 2013; Zhu et al. 2914).

Externally-imposed food availability is difficult to separate from internally controlled processes such as food consumption rate or accumulation and peroxidation of lipids – which in plankton crustaceans are the chief storage nutrient, in addition to general importance for cellular and membrane health. After the initial enthusiasm about lipid peroxidation (LPO) assays as a measure of aging-related oxidative damage and a possible target for antioxidant-based life-extension interventions (Leibovitz and Siegel 1980) the realization came that the use of LPO as a hallmark of aging should be interpreted with caution, particularly when measured by TBARS method of quantifying the final product of LPO, the Malondialdehyde (MDA). It is not clear whether levels of LPO consistently increase with age, why do they, if they do and whether such increase is one of the causes of aging rather than just a concomitated phenomenon (Barja de Quiroga et al. 1992; Praticò 2002), and whether they reflect processes that govern the trade-offs between longevity and oxidative damage-prone processes such as respiration or reproduction (Monaghan et al. 2009). Furthermore, in addition to well-documented protein- and DNA-damaging effects of LPO products through formation of adducts, lipid peroxidation processes are known to function as signaling mechanisms that activate protective cellular functions (Dmitriev and Titov 2010; Ayala et al. 2014). The accumulation of the toxic products of LPO, malondialdehyde (MDA) and 4-Hydroxy-2-Nonenal (4-HNE) is therefore the result of a balance between the availability of polyunsaturated fatty acids (PUFAs) as the primary targets of LPO, utilization of intermediate products of their formation, or utilization of MDA by aldehyde dehydrogenase and glyoxalase pathways (Dmitriev and Titov 2010; Ayala et al. 2014). Thus, instead of being a hallmark of aging and oxidative damage, LPO levels can be a measure of efficiency of PUFA accumulation or antioxidant protection, i.e., processes that may increase, not decrease the lifespan. This is particularly true when LPO is measured not by the accumulation of the toxic final product such as MDA, but by the transient abundance of lipid hydroperoxides themselves, for example by in-situ BODIPY die fluorescence (Pap et al. 1999).

Comparisons of LPO in long-lived and short-lived organisms (or under long- and short life promoting conditions) are rare. Mostly, several analyses have been published correlating lifespan with PUFA content (or integral index of LPO susceptibility, the peroxidation index, PI; Hulbert et al. 2014) rather than directly measured LPO. These studies repeatedly indicated a negative correlation between the PI values and longevity (Munro and Blier 2012; Hulbert et al. 2014; Moghadam et al. 2015) on the context ranging from mild longevity differences between evolved and control *Drosophila* lines and between larval density treatments (Moghadam et al. 2015) to extreme longevity in long-lived bivalves (Munro and Blier 2012).

Data on changes in LPO with age within an organism or a cohort are more abundand. While in humans LPO (measured either directly or as MDA proxy) has been firmly established as a marker of age-related diseases (Miró et al. 2000; Yavuzer et al. 2016; Ramsden et al. 2022), in other organisms the results are more controversial. In *Daphnia,* LPO measured as MDA abundance shows a clear increase with age (Barata et al. 2005, except for the oldest ages; Constantinou et al. 2019; Cai et al. 2020, except that it is not clear how the data were normalized to body weight), while LPO measured as lipid hydroperoxides showed either a moderate increase with age (Cai et al. 2020), or either increase or decrease with age depending on the phase of the ovary cycle during which the measurements are taken (Lowman and Yampolsky 2023), probably reflecting dynamics of PUFAs accumulation in tissues. It would be, therefore, of interest to characterize long- and short-lived *Daphnia* clones using both methods, in the context of short- and long-lived clones.

Thus, the purpose of this study is there-fold. First, we aimed to estimate the amount of variation in lifespan among clones of *Daphnia magna* and among laboratory conditions commonly used in longevity studies in order to increase the likelihood that future interventions studies will be comparable in terms of the baseline (control) longevity. Second, keeping in mind that, arguably, longevity intervention studies aiming at extending uncompromised lifespan are most useful when conducted on genotypes, and under conditions ensuring highest “normal” lifespan in the control group, we aimed to identify genotypes and protocols that allow that. Finally, we aim to characterize the long-lived genotypes with respect to their relation to caloric restriction and to the level lipid peroxidation they are characterized by, across the lifespan.

While meta-analysis across species and different labs may be useful (Cho et al. 2022), there is also an inherent difficulty in such analysis, because of a large number of uncontrolled differences between protocol implemented in different labs, sometimes not fully reported. Here we use a different approach, analyzing only control (untreated) groups from a series of experiments conducted on the same species (*D. magna)* in just two collaborating labs that implement nearly identical protocol differing only in a small number of parameters, such as temperature, medium, group size, and neonate removal method, and using the same set or overlapping sets of reference genotypes (Supplementary Table S1). The disadvantage of this approach, as any other similar meta-analysis, is that it is not, for most types of comparisons, a common garden analysis. Most of the experiments analyzed were designed to test various hypotheses about aging and longevity, such as about effects of maternal age (Anderson et al. 2022b), hypoxia (Ekwudo et al 2022), or biochemical interventions (Pearson and Yampolsky in preparation), or simply to obtain *Daphnia* of different ages for a variety of aging hallmark assays (Anderson et al. 2022a; Lowman and Yampolsky 2023). Hence variation in experimental protocols, in addition to any uncontrolled differences among experiments which may range from quality of algal food to details of maternal treatment or skills of individual investigators. Thus, in addition of analyzing longevity of cohorts from different experiments matched, as much as possible, by a set of conditions, we also report here a more detailed analysis of two common-garden experiments covering 12 geographically distinct isolates of *D. magna* and a range of food levels. Such data would be useful to evaluate natural and lab-generated variation in baseline longevity and to establish a more or less standardized protocol for future studies in this model organism.

## Materials and Methods

### Origin of clones

In this study we jointly analyze 25 longevity experiments (seven of them previously published) conducted in two different laboratories (Supplementary Table S1). Different experiments analyzed in this study included anywhere between 1 and 12 different laboratory clones of *D. magna.* All clones have been obtained from Basel University *Daphnia* stock collection (Switzerland) and are referred to by either full stock ID provided by the stock collection, or, for brevity, by the first two letters of each clone’s ID indicating country of origin (matching Internet country domain two-letter codes). When more than one clone from the same country were used in an experiment, full names are used. Clones were selected to represent a broad range of *D. magna* across Europe to include both mitochondrial clades, both postglacially populated and ancestral refugium areas (Fields et al. 2018), as well as different habitat types (permanent lakes and ponds, summer-dry ponds, and rock-pools). These biogeographic characteristics are not independent from each other. For example, rockpool habitats are overrepresented in the northern, post-glacially populated parts of the range, while summer-dry ponds are typical for the ancestral Mediterranean refugium range. Furthermore, the clones were selected to represent a range of life-histories, to include both short-lived and long-lived clones, as previously characterized (Coggins et al. 2021; Anderson et al. 2022a,b). Again, this life-history feature is not independent form the type of habitat of clone’s origin, as clones originating from rockpools are known to have short lifespan due to higher genetic load in these small populations (Lohr et al. 2014; Lohr and Haag 2015). Clones’ geographic origin, including the original habitat’s hydrology details are listed in Supplementary Table S2 and Supplementary Fig. S1A). All clones have been maintained in Basel stock collection for over 10 years and in one or both labs participating in this study for at least 2 years.

Sixteen of the 25 experiments reviewed were conducted using a subset of 4 clones (FI-FSP1-16-2, GB-EL75-69, HU-K-6, and IL-M1-8; Supplementary Table S1), which will be hereafter referred to as the reference clones. Additional 4 experiments were conducted using these and other clones, making the total count of datasets that can be used for the 4 reference clones 20 out of the total of 25 datasets. See Results and Supplementary data for the information of which analyzes were conducted using individual clones, the 4 reference clones, or all clones analyzed. Finally, one of the experiments (experiment 25; Coggins et al. 2021) was conducted using mixed cohorts, in which although individuals’ clonal identity was preserved, the number of replicates per clone per treatment was low, resulting in a clone-aware analysis being impossible. This data set was included into the analysis either with all clones combined, or, as appropriate, with only the 4 reference clones included, as appropriate.

### Experimental conditions

*Daphnia* stocks were maintained and longevity experiments conducted in either COMBO (Kilham et al. 1998) or ADaM (Klüttgen et al. 1994.) zooplankton media (reconstructed pond water) at a density of 1 adult female per 20 mL of media, with the media replaced every 3 to 4 days. In the few experiments that included male cohorts male density was 1 adult per either 5 or 10 mL of medium, as males are about half the linear size of adult females. Because clones of *Daphnia* differ substantially in the frequency of males produced under typical laboratory conditions, most experiments included female only, or, if they did include male cohorts, these cohorts were limited to a small subset of clones.

Unless otherwise indicated, *Daphnia* were fed with a suspension of *Scenedesmus acutus* (synonym: *Tetradesmus obliquus*) green algae grown in the lab on constant light in one of commonly used phytoplankton grown media (typically either Kilham et al. 1998, or Jüttner et al. 1983) with vitamins cocktail added as described in Goulden et al. (1982). Majority of experiments included in the present analysis have been conducted under the feeding regime consisting of daily addition of the algal suspension to the concentration of 10^5^ cell/mL, which corresponds to 3.6 μg C mL^-1^ day^-1^, a baseline concentration chosen as being slightly below complete consumption of food provided within 24 hours, and thus corresponding to a mild dietary restriction (see below). Algae concentration was measured by means of far red fluorescence with blue light excitation using Qubit fluorometer (Thermo Fisher) calibrated by direct cell counts in a hemocytometer. Several experiments included treatments with food concentrations 2-4 times lower or 2-4 times higher than the “standard” concentration, allowing the analysis of starvation and caloric restriction effects on longevity. Three of the experiments included, in addition to the regular diet, feeding treatments with the diet consisting of either a stramenopile alga *Nannochloropsis,* with the concentration matched to the *Scenedesmus* diet by carbon content, or dry and resuspended cyanobacterium *Spirulina sp.*, matched to the *Scenedesmus* diet by dry weight. Details of feeding regimes in each experiment are listed in Supplementary Table S1. Further details of experimental procedures of previously published experiments (Supplementary Table S1) are available from the corresponding publications (Coggins et al. 2021; Cho et al. 2022; Ekwudo et al. 2022; Anderson et al. 2022a,b; Lowman and Yampolsky 2023; Moore et al. 2023).

### Cohorts with individual housing

In experimental cohorts in which *Daphnia* were maintained individually *Daphnia* were housed in 35 mL glass shell vials containing 20 mL of medium. Water was changed and neonates removed within 24 hours after the release of each brood of neonates or after a molt resulting in no neonates produced (i.e., every 3-4 days), minimizing effects of competition between the mother and the offspring for food.

### Cohorts with small group housing

In most of the lifespan experiments analyzed in this study *Daphnia* cohorts were maintained in either groups of 5 in 100 mL glass jars or in groups of 10 in 200 mL jars (females and males separately; male’s density as described above). When mortality occurred (or individuals were removed from cohorts for various measurements) water volume in each jar was adjusted according to the number of remaining individuals; jars sharing the clone and data of birth were combined when there were fewer than 6 or 11 individuals remaining in two such jars, in 100-mL jar and 200-mL jar experiments, respectively. Thus the group size in this treatment was either 5 or 10 at the start of the experiment and below these values as the cohorts were dying out. It also results in individual jars not being treated as independent replicates in longevity analysis.

Both the 100-mL jar 200-mL set-ups can be run with manual handling of individuals at the time of water change or in the semi-manual version with *Daphnia* housed within 50 nL plastic inserts that can be moved to fresh jars, simultaneously removing any neonates present in the jar. To accomplish this the inserts are equipped with 1 mm nylon mesh that retain the adults but allow neonates to flow through. In order to prevent inadvertent introduction of offspring into the cohort resulting in potential “Jeanne Calment effect” (Milholland 2020) due to incomplete filtering out of neonates by the insert mesh, all inserts were inspected daily for any retained juveniles. The use of inserts, therefore, allows to eliminate manual handling of individuals and automatic removal of neonates during or between water changes. Removal of neonates occurred in different experiments that utilized the inserts either every other day, or once in 4 days simultaneously with water change.

### Cohorts with large group housing, including “Smart Tanks”

In three of the experiments *Daphnia* cohorts were maintained in larger groups (50 individuals in 1L containers initial cohort set up), resulting in fewer replicate containers. Details of this experimental set up (“Smart Tanks” set up), including the automatic removal of neonates from the experimental containers and semi-automatic census of cohorts’ size have been published elsewhere (Cho et al. 2022).

### Feeding rate and lipid peroxidation and lipofuscins measurements

In the experiment involving 12 different clones that aimed to test hypotheses about physiological correlates of clone-specific longevity (Experiment 18, Supplementary Table S1) we measured feeding (filtration) rate in *Daphnia* of different age, as previously described (Anderson et al. 2022a). Briefly, individual *Daphnia* were placed, in 3-4 replicates per clone, in 1.5 mL vials containing zooplankton medium with the initial *Scenedesmus* algae added to the concentration of 2×10^5^ cells/mL and algal concentration was measured by Qubit fluorometer 1.5, 3, and 4.5 hours after the start of the experiment in both *Daphnia-*containing and blank vials. The slope of blank-subtracted linear regression over time was used as a measure of feeding rate, converted from fluorescence units to cell concentration using the standard calibration, and normalized by *Daphnia* wet weight.

In the same experiment we also measured abundance of lipofuscins, hydroperoxidized lipids (LPO) and the accumulation of malondialdehyde (MDA), the secondary product of lipid peroxidation. Lipofuscins abundance was measured in subset of 9 out of 12 *Daphnia* clones at ages 25 and 55 days by autofluorescence as described in Lowman and Yampolsky (2023). Whole body fluorescence was measured excluding the gut and the highly mandibles which are highly autofluorescent regardless of the age. Lipid hydroperoxides were quantified by fluorescence microscopy of 4 body regions – the spiral section of the nephridium, 2^nd^ epipodite (a thoracal leg ramus with excretory functions), abdominal fat body, and the ovary, stained with BODIPY C11 dye (Pap et al. 1999), as previously described (Anderson et al. 2022a; Lowman and Yampolsky 2023). MDA concentration in whole-body extracts was measured by TBARS method using Caiman Chem (Ann Arbor, MI, USA) kit, as described in Coggins et al. 2017).

### Joint analysis across experiments

We jointly analyzed a number of published and unpublished experiments to evaluate the longevity differences between environmental conditions (such as temperature or group size) and across labs and establish means and variances of the baseline longevity. These experiments shared the density at which female *Daphnia* were maintained (1 individual per 20 mL) and, most of them, food type and concentration used (*Scenedesmus,* 100,000 cells/ml/day per individual), but differed in *D.magna* clones used, group size in which the cohorts were maintained (from individually to groups of >100 in a single “Smart Tank” insert, Cho et al. 2022), medium used (COMBO vs. ADaM reconstructed pond water media), temperature at which they were conducted, location of the experiments (HMS vs. ETSU), the personnel accomplishing censuses and water changes, and whether the neonates were removed manually, semi-manually, or through the action of the “Smart Tank” set up. Not all these factors could be analyzed separately, as some of the factors, such as laboratory and temperature and clones used are confounded. Additionally, we included two experiments in which a different food source was used (*Nannochloropsis* alga) and two experiments in which male cohorts were analyzes alongside the standard female-only cohorts. Many of these experiments had additional treatments, such as food concentration or addition of a potential metabolism-modifying agent. In such cases only the data from control cohorts adhering to the standard conditions were used. The survival data from these experiments were not analyzed together in a single survival analysis model. Rather, each experiment yielded median lifespan (50-th percentile survival time), as well as 25-th and 75-th percentile survival times, and these aggregate values were analyzed together. Each independent cohort (e.g., initiated on a separate date) within each experiment yielded one set of survival quantiles.

Because we observed the effect of group size on longevity that was independent of density and food supply, we tested the hypothesis that the increased mortality in *Daphnia* maintained in groups is caused by clusterization of deaths within the same replicate jar and the same 48-hour observation period. To do so we observed the number of death occurring as single events to those that occurred in clusters of 2-5 in *Daphnia* maintained in groups of 5 individuals per 100 mL of medium, as well as in clusters of 2-4 in the ones maintained in groups of 4 individuals per 80 mL. These 2 data sets came from the same 2 experiments (#18 and 20), with 5 individuals per 100 mL jar treatment, and 4 individuals per jar remaining after a single mortality event. These counts were compared to prediction based on binomial probabilities calculated on the basis of overall probability of death per observation period in such groups, with the number of individuals in a group (5 or 4) as the number of trials, and death cluster size as the number of successes. As the expected counts of events with 4 or 5 deaths occurring together are low contingency analysis including frequencies of such evens is biased. Therefore we compared singleton death count to clustered death count pooled together using Fisher’s Exact Test.

Statistical analysis has been performed using model fitting and survival analysis platforms of JMP (Ver. 17; SAS Institute 2018). Specifically, for survival analysis Kaplan-Meier method was used for the estimation of mean and median lifespan in a single-factor analyses and Proportional hazards method for multi-factor models.

Because the outcomes of statistical tests that involve age as a factor can be radically affected by the subset of individuals surviving to each age, each analysis has been conducted in four versions: retaining all absolute ages and all clones, retaining all relative ages (normalized by clone-specific median lifespan), retaining a subset of clones surviving to all ages, and retaining a subset of ages to which all clones have survive and thus could be measured. In no case reported below any substantial difference in outcomes between these four analyses has been detected and, unless otherwise indicated, only the results of the latter approach are reported.

## Results

### Correlations with clone’s habitat of origin

Clones radically differed in lifespan across a variety of experiments and experimental conditions (Fig. 1 – 4; Tables 1, 3). As predicted, clone-specific median lifespan decreased along the north-south gradient, stronger in lower food treatments than in high food treatments (Supplementary Fig. S1, A,B). This decrease was not very well supported given the low number of clones (P<0.02), but substantial in magnitude, with northernmost clones showing nearly 2-fold reduction in lifespan relatives to the southern-most ones. Unexpectedly, a similar cline was observed in a longitudinal comparison (Supplementary Fig. S1, C), probably largely due to a slight SW-to-NE spread of clones’ origin locations. The comparison between habitat types, showed, as expected, a higher median lifespan in clones from permanent habitats than in those originating from rockpools, although none of the comparisons was statistically significant (Supplementary Fig. S1, E-G). It was not possible to untangle the potential effects of latitude and habitat types due to a strong confounding between the two parameters. Either way, geography has a weaker effect on clone-specific longevity than other clone-specific parameters (see below).

**Fig. 1.**
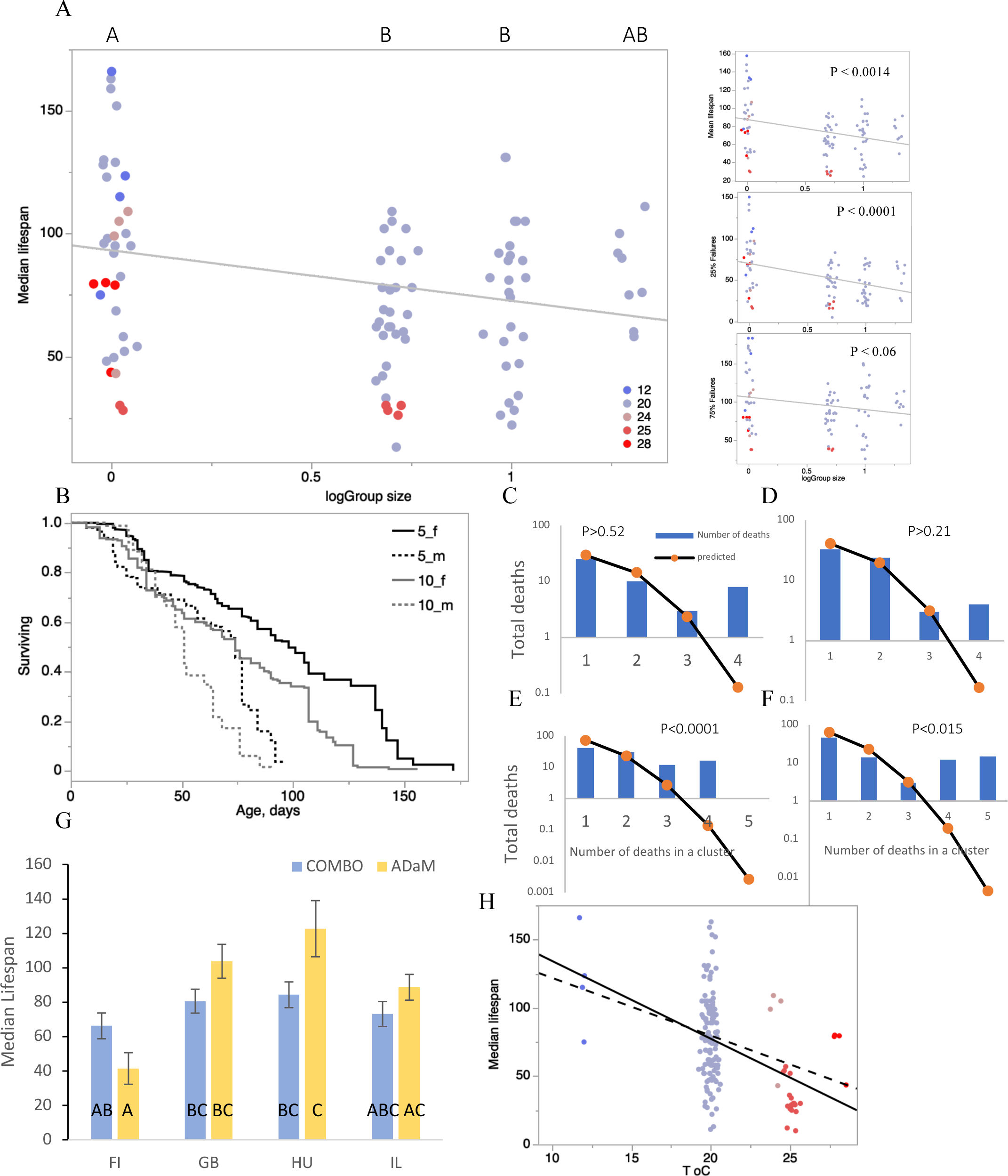
Effect of group size (A - F), medium composition (G) and temperature (H) on lifespan of *Daphnia* cohorts in the 4 reference clones. A: Lifespan measured in various experiments with different group size, shown as log_10_. Colors indicate temperatures between 12 and 28 °C (blue to red). Regression line based on 20 °C experiments only; others shown for comparison. Letters on top indicate Tukey test of the effect of group size on lifespan at 20 °C, if group size treated as a categorical variable; categories not sharing a letter are different at P<0.05. The same Tukey test results apply to median, mean and 25% quartile survival. No categories are different from each other for the 25% quartile survival. See Table 1 for complete statistics. See Supplementary Fig. S1A for the same data for all clones combined across different experiments. B: Survival curves of several cohorts of females (solid lines) and males (dotted lines) of the “IL” clone maintained in either groups of 5 (black lines) or 10 individuals (gray lines). See Table 2 for complete statistics. See Supplementary Fig. S1B,C for the same data with each cohort shown separately. C – F: clustered death analysis. C and E: 12-clones experiment (#18); D and F: caloric restriction experiment (#20), food level 100. C and D: clumping of death events within 48 h observation period in *Daphnia* maintained in groups of 4 in 80 mL of medium. E and F: clumping of death events in *Daphnia* maintained in groups of 5 in 100 mL of medium. Bars: observed counts of death occurring in clusters of each size. Lines: expected based on Binomial assumption of independence. P-values shown are from 2-tailed Fisher Exact Test comparing the likelihood of deaths to occur in clusters vs. in single events, relative to binomial expectations. G: Median lifespan of females from the 4 reference clones maintained in two media. No individual comparison of the medium composition effect tests survived Tukey test. H: Data across all experiments. Solid regression line for all clones, dashed line for the 4 reference clones only. Colors as on panel A.

**Table 1.** ANOVA of the effects of group size and water composition on lifespan quantiles in the 4 reference *Daphnia* clones. P-values < 0.01 shown in bold.

| Source | DF | 25% quartile |  |  | Median lifespan |  |  | 75% quartile |  |  |
| --- | --- | --- | --- | --- | --- | --- | --- | --- | --- | --- |
|  |  | SS | F Ratio | Prob > F | SS | F Ratio | Prob > F | SS | F Ratio | Prob > F |
| Clone | 3 | 8186.59 | 6.9 | **0.0004** | 20477.2 | 10.1263 | **<0.0001** | 21859.3 | 12.99 | **<0.0001** |
| Group size | 1 | 6430.6 | 16.2 | **0.0001** | 1833.8 | 2.72 | 0.1 | 1046.8 | 1.87 | 0.18 |
| Clone*Group size | 3 | 4048.4 | 3.4 | 0.0225 | 1044.4 | 0.52 | 0.67 | 179.3 | 0.11 | 0.96 |
| Water | 1 | 270 | 0.7 | 0.4 | 2576 | 3.82 | 0.05 | 3725.4 | 6.64 | 0.0125 |
| Clone*Water | 3 | 1379.4 | 1.2 | 0.3 | 8476.5 | 4.19 | **0.009** | 14810 | 8.8 | **<0.0001** |
| Group size*Water | 1 | 4308.2 | 10.9 | **0.0016** | 1763.4 | 2.62 | 0.11 | 1394.6 | 2.49 | 0.12 |
| Clone*Group size*Water | 3 | 1943.1 | 1.6 | 0.2 | 403.6 | 0.2 | 0.9 | 1054.4 | 0.63 | 0.6 |
| Error | 67 | 26556.1 |  |  | 44487.9 |  |  | 35348.6 |  |  |

**Table 2.** Likelihood ratio proportional hazards analysis of the effects of group size and sex on longevity. Individual cohorts nested within group size treatment. Cf. Fig. 1B.

| Source | DF | L-R $\chi^2$ | Prob> $\chi^2$ |
| --- | --- | --- | --- |
| Sex | 1 | 101.6 | <0.0001 |
| Group size | 1 | 45.59 | <0.0001 |
| Cohort [Group size] | 2 | 1.85 | 0.4 |
| Group size*Sex | 1 | 1.36 | 0.24 |
| Cohort*Sex[Group size] | 2 | 2.54 | 0.28 |

**Table 3.**
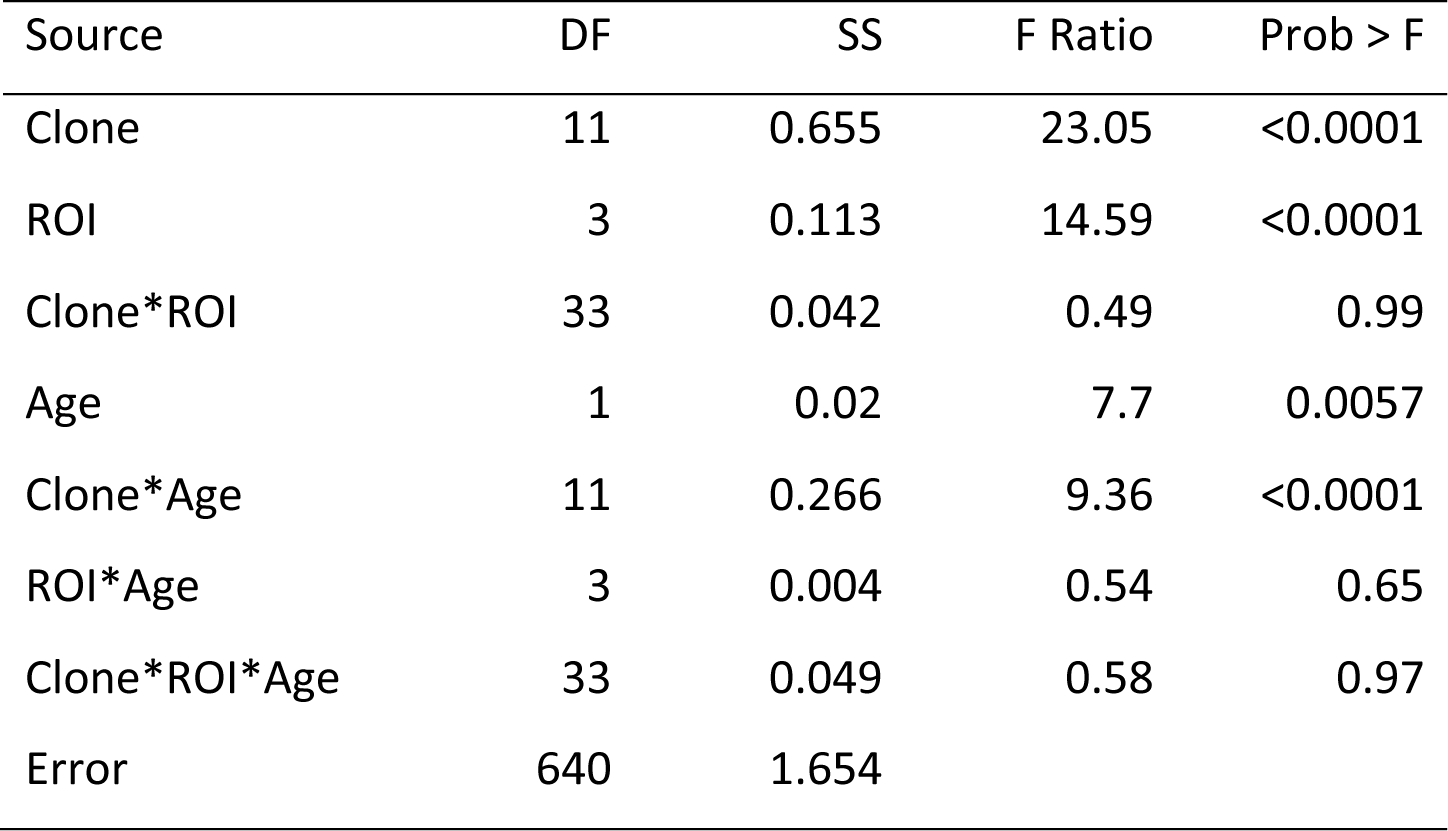
ANOVA of the effects of age, clones, and body regions (ROI) on lipid peroxidation measured by fluorescence microscopy of *Daphnia* stained with BODIPY C11 dye. Age is treated as a continuous covariable. P-values <0.0001 shown in bold. See Supplementary Table S7 for variants of the same analysis with normalized age and with short-lived clones not measured at old ages excluded. See Supplementary Table S8 for variants of the same analysis for each ROI separately.

| Source | DF | SS | F Ratio | Prob > F |
| --- | --- | --- | --- | --- |
| Clone | 11 | 0.655 | 23.05 | <0.0001 |
| ROI | 3 | 0.113 | 14.59 | <0.0001 |
| Clone*ROI | 33 | 0.042 | 0.49 | 0.99 |
| Age | 1 | 0.02 | 7.7 | 0.0057 |
| Clone*Age | 11 | 0.266 | 9.36 | <0.0001 |
| ROI*Age | 3 | 0.004 | 0.54 | 0.65 |
| Clone*ROI*Age | 33 | 0.049 | 0.58 | 0.97 |
| Error | 640 | 1.654 |  |  |

**Table 4.**
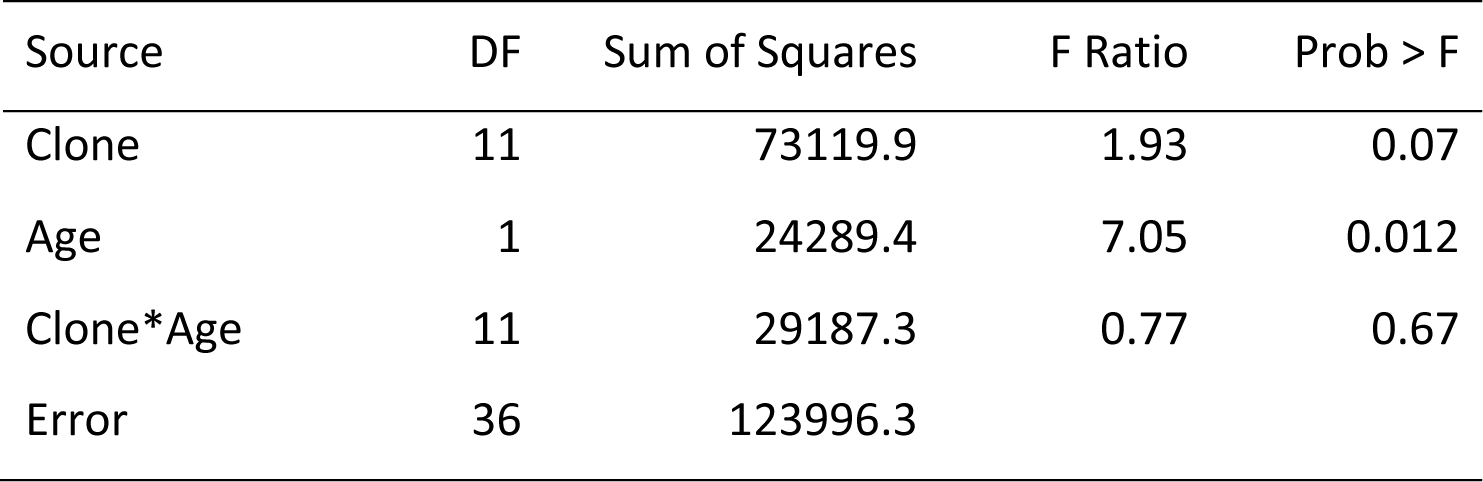
ANOVA of the effects of age and clones on the amounts of lipid peroxidation secondary product MDA (µM/mg protein) measured in whole body extractions. Age is treated as a continuous covariable.

### Effects of group size, medium, and temperature on cohorts’ longevity

Across a number of experiments that differed in group size and medium, but shared other experimental conditions, there was a moderate in magnitude, but statistically significant effect of group size, with individually housed *Daphnia* showing the highest longevity measured as either median or mean lifespan (Fig. 1A, Table 1, Supplementary Fig. 2 A). The effect was the strongest for the 25%-quantile of the lifespan and nearly non-significant for the 75%-quantile, indicating that larger group size contributes to early mortality. The effect of group size was observed in both females and males with a similar magnitude as reflected by no significant sex-by-group size interaction in the proportional hazards analysis (Fig. 1B, Table 2). This indicates that it is unlikely that group size affects female mortality by increased exposure to neonates produced in the course of the lifespan experiments.

The effect of group size manifested itself in the form of clustered deaths (coinciding within the same jar within a single 48-h census period), which occurred more frequently than expected by chance when *Daphnia* are maintained in groups of 5 individuals (Fig. 1 E,F). This overrepresentation of clustered deaths is due almost entirely to events in which all or all but one individuals in a group suffered mortality. In groups of 4 individuals per 80 mL of medium, however, the enrichment in clustered deaths is no longer observed. This clusterization of deaths might be indicative of either infections or emission of germs of toxins from carcasses shortly after death. However, zero mortality was observed (in both controls or treatments) in 5-day trials with *Daphnia* of similar age being exposed to either water from the jar in which clustered deaths occurred, or carcasses of individuals who died in a clustered manner, or homogenized such carcasses, as well as to carcasses of individuals who dies in sporadic manner (data not reported). This indicates that whatever causes clustered mortality is either short-lived, or volatile, or attached or absorbed to the surface of the jar, or entirely extraneous such as a handling error.

The use of meshed jar inserts (semi-manual neonate removal method) resulting in no significant effect on longevity relative to no inserts used (manual neonate removal), in either 100 or 200 mL jars (comparison based on 4 and 2 clones, respectively; Supplementary Fig. 2 B, Tukey test P > 0.05), although in 5 out of 6 comparisons *Daphnia* lived slightly longer when inserts were used.

The comparison of the water composition (COMBO vs. ADaM media) in the same experiment revealed a mild increase in longevity (particularly in the 75% survival quantile age) in the ADaM medium (Fig. 1G, Table 1). However, there was also evidence on clone-by-medium interactions and none of individual comparisons between water compositions within the four reference clones survived post-hoc Tukey test.

Predictably, lifespan radically decreased with temperature (Fig. 1H). Full factorial analysis was possible only for 2 of the clones (GB-EL75-69 and IL-M1-8) represented in all temperatures, the analyses including all clones or the 4 reference clones could not estimate all parameters of the linear model. However, the clone-by-temperature parameter could be estimated in all cases and it was never significant, regardless of which set of clones was included into the analysis (data not presented).

### Between-labs and within-lab-between-experiments comparisons

Only 2 lifespan experiments conducted in two different laboratories (Harvard and ETSU) could be matched by food, temperature, group size conditions and by the clone used. Fig 2 shows a remarkable agreement between the 2 independent experiments in both female (Fig. 2A) and male (Fig. 2B) cohorts. On the other hand, two nearly identical experiments conducted in the same lab (ETSU) by two different investigators with a significant time overlap showed consistent differences in estimated lifespan duration (Fig. 2 C, D). The only controlled difference between the two experiment was that in one of them (Experiment 18) neonates were removed every other day, while in the other (Experiment 20) once in 4 days.

**Fig. 2.**
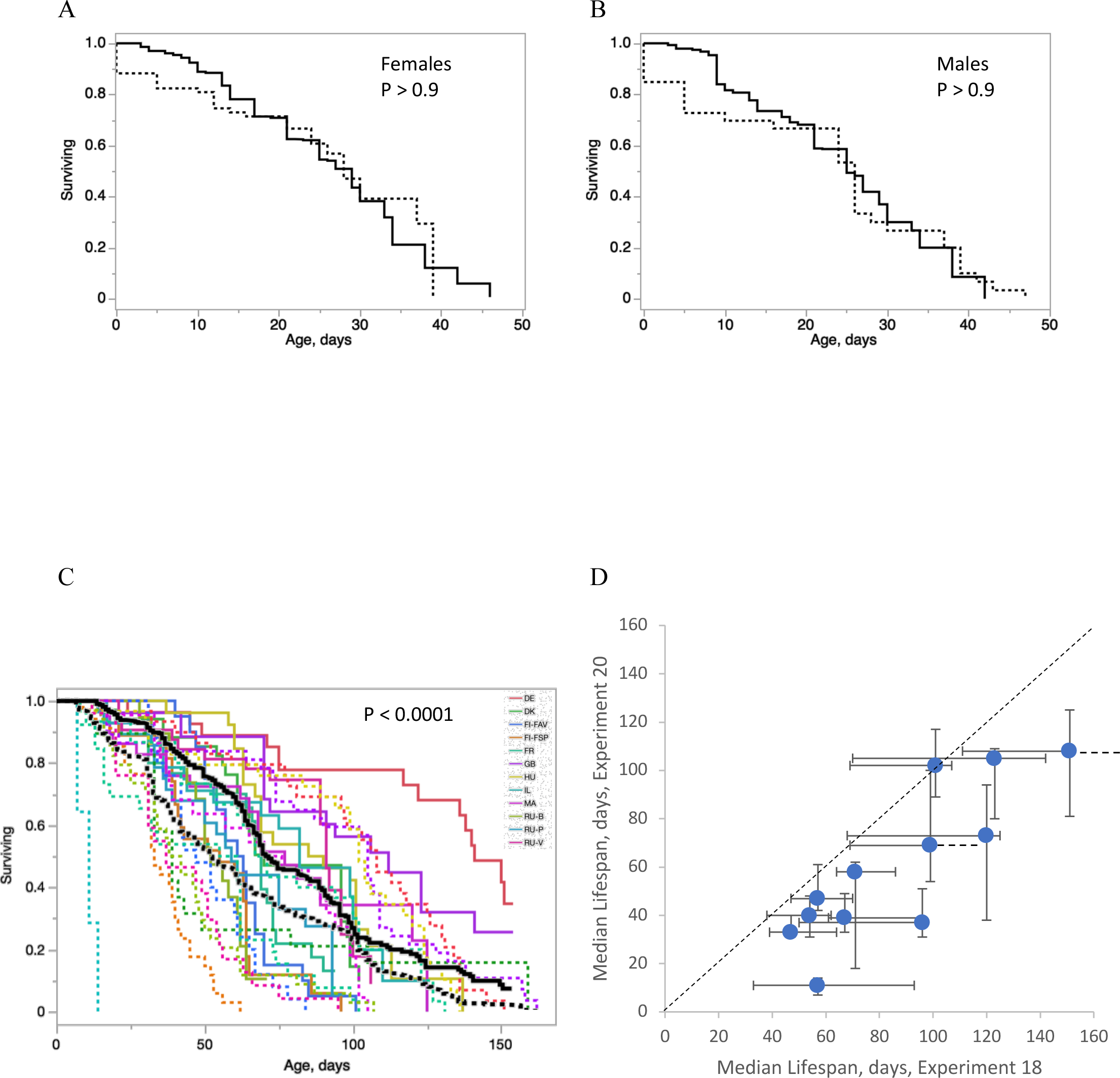
Comparison across labs (A,B) and between comparable experiments within a lab (C,D). P-values from log-rank test between the two labs or between two experiments. A, B: Solid line, Harvard Medical School; dotted lines, East Tennessee State University. Two experimental cohorts match by food, group size (5), temperature (25 °C), and *Daphnia* clone used (“IL”). A: female cohorts; B: male cohorts. See Supplementary Fig. 1D for the same data with each cohort shown separately. C, D: two identical experiments (12 reference clones, groups of 5, 20 °C) conducted simultaneously in the same lab (ETSU) by two different investigators. C: Survival curves with colors indicating 12 different clones; black lines, overall survival; solid lines, Experiment 18, neonates removed every other day; dotted lines, Experiment 20, neonates removed once in 4 days (same data as on Fig. 4A, middle panel, see below). D: Median lifespan comparison across clones. Bars represent 95% confidence intervals (dashed lines where upper CI estimation was not possible). Rank correlation over clones: π = 0.85. Diagonal dashed line represents the equal lifespans expectation.

### Sex differences in lifespan

Across several experiments conducted at different temperatures and with different group sizes common garden male cohorts showed slightly reduced lifespan relative to their female counterparts (Fig. 3), with null-intercepted regression line showing the slope of 0.774. This difference was not universal though. While clearly aging differently in experiments conducted at 20 °C Fig. 1B, Supplementary Fig. S2 C,D, Table 2), males and females showed similar median lifespan at 25 °C (Fig. 2, Supplementary Fig. S2 E), and in one case a male cohort survived significantly longer than the corresponding male cohort. Males’ lifespan also responds differently from that of females to food concentrations at what is below *ad libitum* for females (Fig. 2 B,C; Supplementary Table 3). XXXX

**Fig. 3.**
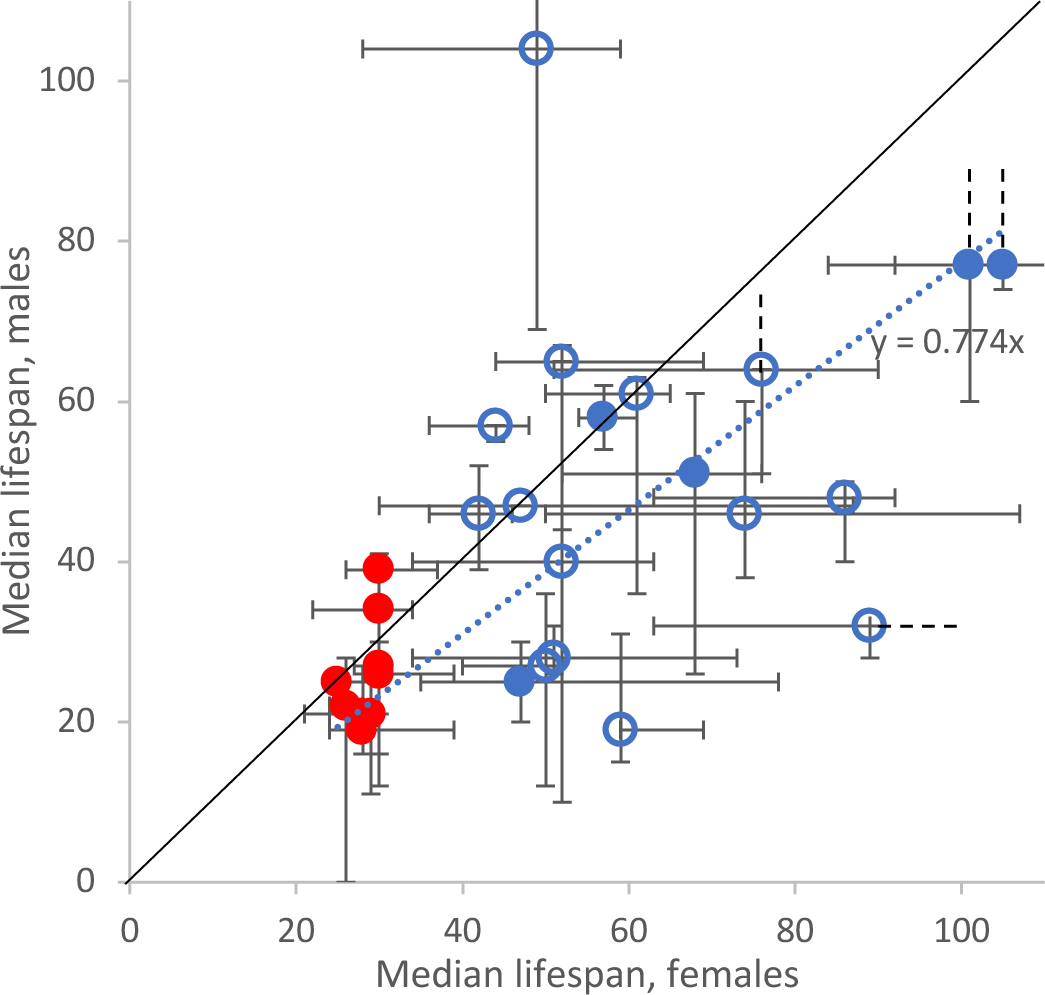
Males vs. females median lifespan measured in the 4 reference clones at two temperatures (24°C, red or 20°C, blue) and under two group size regimes (5, filled symbols, or 10, open symbols individuals in a group, 20 mL of medium per individual and 1×10^5^ cells/mL/day food level in both). Bars represent 95% confidence intervals (dashed lines where upper CI estimation was not possible). Regression line drawn through intercept=0; regression coefficient r = 0.755 ± 0.046 (SE); P<0.0001. Diagonal dashed line represents the equal lifespans expectation.

### Effects of caloric restriction

The above data was collected at food concentrations below the *ad libitum* level for females. The inclusion of higher food levels (above 2 ×10^5^ *Scenedesmus* cells/mL/ day) revealed that the median lifespan did not change within the range of food levels between 0.5×10^5^ and 3×10^5^ *Scenedesmus* cells per mL per day and dropped below and above these levels (Fig. 4, Supplementary Fig. S3). In most clones the “standard” food level of 1×10^5^ cells/mL/day resulted in the highest median lifespan, but in few clones there was a significant differences between 0.5×10^5^ and 1×10^5^ food levels (Fig. 4A, B, C). The 4-fold increase in food concentration to the above *ad libitum* level of 4×10^5^ cells / mL / day resulted in a significant (∼30%) reduction of lifespan in the long-lived genotypes but causes no difference in lifespan of short-lived ones (Fig. 4D). Note that Fig. 4D shows the comparison of caloric restriction effect and baseline lifespan in restricted food treatment obtained in two separate independent experiments, i.e. the observed correlation is not an autocorrelation artifact. The *ad hoc* classification of the clones into long-lived, caloric restriction-responsive and short-lived, caloric restriction non-responsive ones becomes apparent again in the analysis of the effects of lipid peroxidation on lifespan (see below). Further analysis of caloric restriction on lifespan, including the relationship between reproduction rate and longevity at different food levels will be reported elsewhere (Bright and Yampolsky, in preparation).

**Fig. 4.**
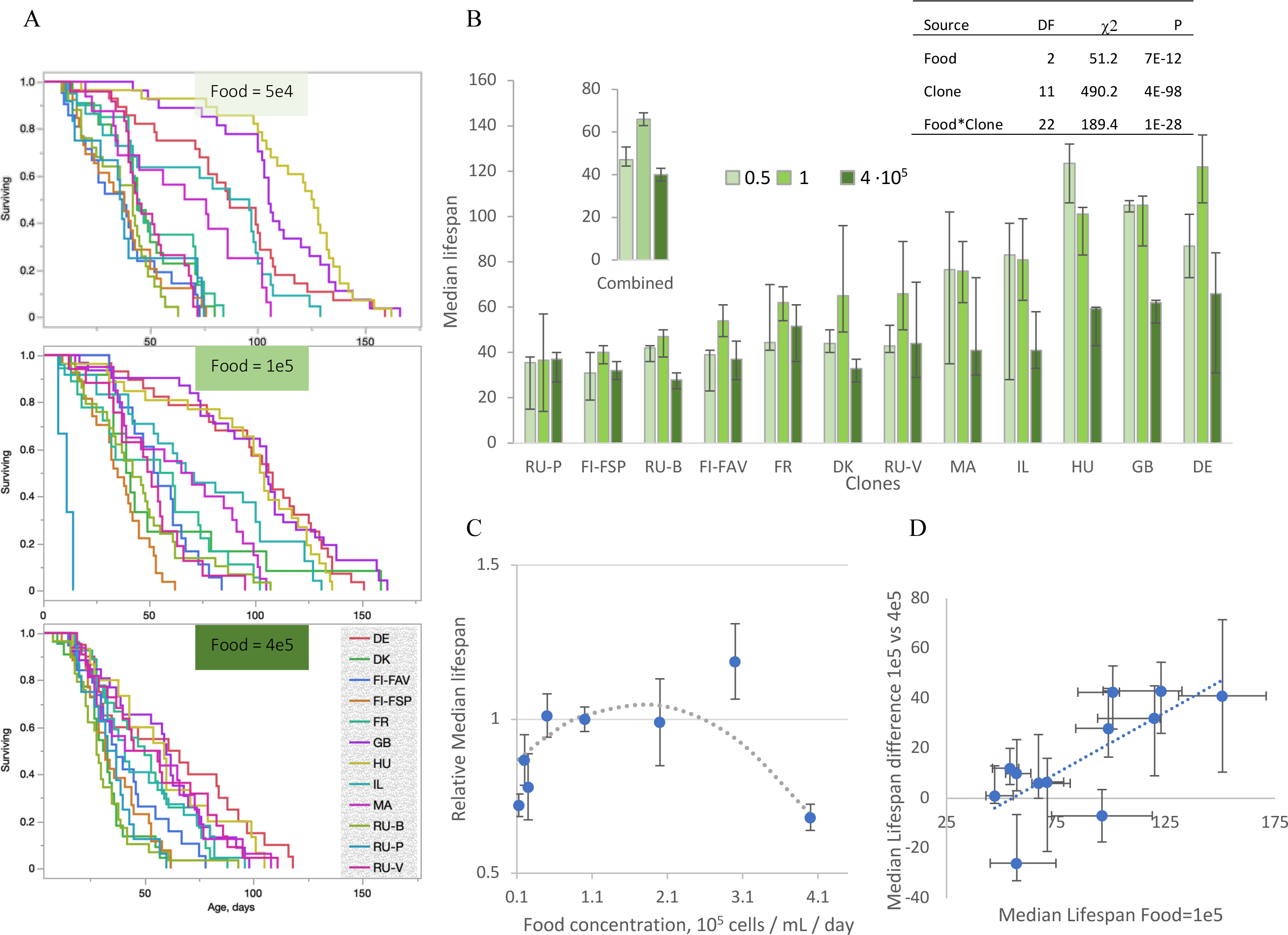
Effects of caloric restriction on *Daphnia* lifespan. A: Survival curves of 12 clones of *D.magna* under *ad libitum* level (4 10^5^ algal cells / mL / day, bottom), 1 10^5^ algal cells / mL / day (middle) and 5 10^5^ algal cells / mL / day (top). B: The same data represented as clone-specific median lifespan at 3 food levels. Abbreviates clone IDs as in Supplementary Table S2. Vertical bars are 95% confidence intervals. Inserts: the same data, 12 clones combined and a proportional hazard analysis of the clonal and food differences; note a significant clones-by-food level interaction. 1 10^5^ algal cells / mL / day treatment: combined data from two experiments. C: combined data from several lifespan experiments (Supplementary Table S1) that included different food levels conducted in a subset of 4 clones. Vertical bars are SEs. See Supplementary Fig. S2 for similar data for 2 reference clones maintained at 5 food levels. D: correlation between the strength of caloric restriction effect measured as the clone-specific difference in median lifespan between *ad libitum* and restricted food treatments (1 10^5^ and 4 10^5^ algal cells / mL / day, respectively) and clone-specific median lifespan at restricted food measured in an independent experiment. Bars are SEs. A, B: [[ExpID=18, 20]] C: [[ExpID=18, 20]] D: [[ExpID=1, 6, 13, 18, 19, 20, 23, 24]]

### Effects of food type

Only 2 experiments in our dataset utilized both food species, *S. acutus* and *N. limnetica* in a common garden set-up (experiments 24 and 25, Supplementary Table S1; Coggins et al. 2021; Moore et al. 2023, respectively) and one experiment compared the standard *S. acutus* diet to a dry weight-matched diet consisting of *Spirulina*. Different *N.limnetica* experiments differed in group size and temperature, precluding a joined analysis. In each experiment analyzed separately, the *Nannochloropsis-*fed *Daphnia* showed slightly extended lifespan at 25 °C, when all clones were considered (proportional hazards P<0.006; Coggins et al. 2021), but showed no difference in lifespan from the *Scenedesmus*-fed *Daphnia* from the 4 reference clones analyzed separately (P>0.75). In the other experiment conducted at 20 °C on the same 4 reference clones the *Nannochloropsis* diet extended the lifespan of 2 short-lived clones and shortened the lifespan of the two long-lived ones (proportional hazards clone-by-food interaction term P<0.0001; Moore et al. 2023). This indicates that the effects of the two types of diets on the lifespan are highly genotype-specific. A common garden comparison between longevity of one clone (IL) on freshly grown *S. acutus* vs. dry and resuspended *Spirulina* diet revealed a significant reduction of lifespan on *Spirulina:* 61 vs. 29 days median lifespan, respectively (13-51 vs. 38-70 days 95% CI; Proportional Hazards loglikelihood ratio test P < 0.0063; data not presented).

### Lack of trade-offs between fecundity and longevity at calorically restricted food level

In the experiment that involved 12 clones as was conducted at 1 10^5^ cells/mL/day food level fecundity decreased with age (Supplementary Fig. S4 Supplementary Table S4), dropping from the average of approximately 1.5 neonate per female per day (i.e. 4-6 eggs per clutch to approximately 0.75 neonate per female per day (i.e. mean clutch size between 2 and 3; these averages include skipped clutches where no neonates are produced in a molting cycle). Age, clone, and clone-by-age interaction effects were all significant. Neither overall fecundity with all ages combined, nor age-specific fecundity within each 1-day age class showed any trade-off (e.g., negative correlation) with clones’ median lifespan (Supplementary Fig. S5). In contrast, all regression were positive, and for the mid-age classes between 30 and 70 days significantly so.

### Age-related changes in feeding rate, lipofuscin accumulation and lipid peroxidation and their effect on lifespan

The above analyses were concerned with clonal or experimental parameters that did not change throughout the life of an experimental cohort. In addition to this we analyzed 3 parameters we expected to change over the lifetime of cohorts and their correlations with lifespan: feeding rate and two measures of lipid peroxidation. Feeding rate significantly decreased with age (Supplementary Table S5; Supplementary Fig. S6 A), regardless of whether it was normalized by *Daphnia* wet weight, and regardless of whether all clones and all ages were included into the analysis, or only those ages in which all clones were represented. Clones showed a significant variation for weight-normalized feeding rate, but not absolute feeding rate, indicating that food consumption per individual was similar in all clones. There were no clone-by-age interaction in any of the tests and the rate at which feeding rate decreased with age, correspondingly, showed no correlation with clone-specific medial lifespan measured in the same experiment (Supplementary Fig. S6 B), although there is a slight tendency of a negative correlation, particularly among the long-lived, CR-responsive clones: clones with the longest lifespan decreases their feeding rate with age faster than those with the shortest lifespan, with 2 outlier clones that eliminate the significance of the pattern (Supplementary Fig. S6 B). Overall, there was a positive correlation between longevity (mean and median lifespan) and early life feeding rate (Fig. 5A), but not between longevity and mid-life feeding rate (Fig. 5B).

**Fig. 5.**
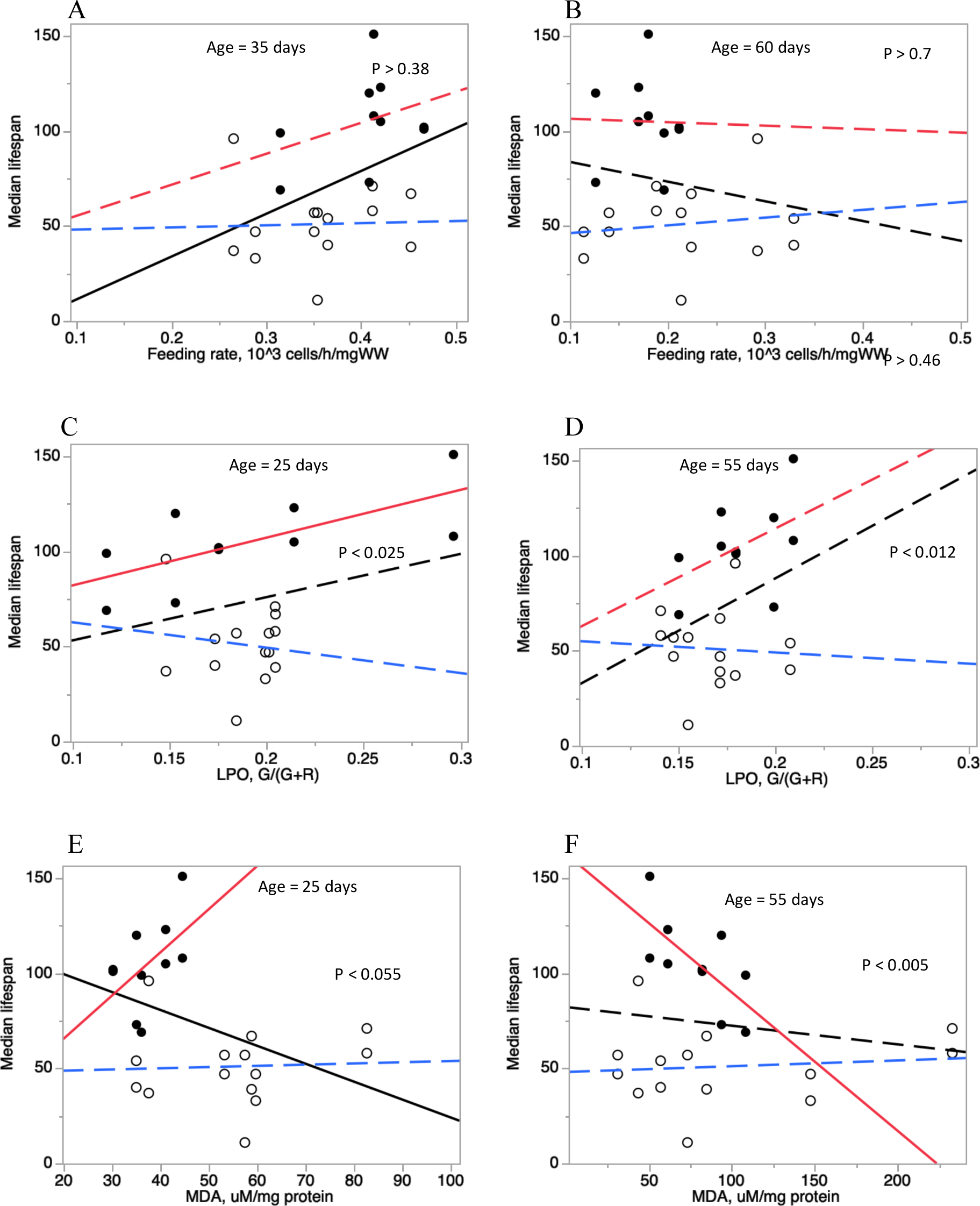
Clone-specific median lifespan (measured in two independent experiments) plotted against feeding rate (thousands of cells consumed per hour per mg wet weight, A, B), LPO measured by in-vivo BODIPY fluorescence (ratio of green fluorescence to the sum of green and red fluorescence; average of 4 tissues; C, D), and LPO measured by whole-body extract MDA concentration (µM / mg protein; E, F). Each of the three parameters measured in young individuals (age 25-35 days, A, C, E) and mid-life individuals (age 55-60 days, B, D, F). Linear regressions shown for all clones (black lines) and separately for the long-lived, caloric restriction sensitive and short-lived, caloric restriction insensitive clones (black and open symbols and red and blue regression lines, respectively). Regression lines with P>0.05 shown as dashed lines. P-values represent the significance of the interaction term between the clone type and the covariable (heterogeneity of slopes of red and blue regression lines). See Supplementary Table S9 for full statistical analysis and Supplementary Fig. S8 for the data shown here on parts C and D, with each tissue analyzed separately.

Lipofuscin abundance increased significantly between the age of 25 and 55 days, with mean background-subtracted autofluorescence increasing from 2.391 (SD=1.222) to 3.541 (SD=1.618), with a highly significant difference among clones, but no evidence of clone-by-age interaction (Supplementary Table S6, Supplementary Fig. S6 C, D). The amount of fluorescence increased through the increase of the number of spots, the size of spots and the intensity within spots (data not presented). Unexpectedly, the amount of increase in lipofuscins between ages 25 and 55 days was weakly positively correlated with clone-specific lifespan (Supplementary Fig. S6 D).

The two LPO parameters measured, BODIPY fluorescence in in-vivo stained tissues and MDA concentration in whole-body extracts behaved differently with age (Supplementary Fig. S6 E, G). BODIPY fluorescence showed a slight decrease with age (Supplementary Figs. S6 E). The age effect was slight, but significant regardless in all types of analysis (absolute vs. relative age, all ages vs. only ages in which all clones were represented, all clones vs. only clones surviving to the highest age at which measurements were done, Supplementary Table S7). Likewise, there were significant differences among clone and among tissues studies, with no interaction between the two. When tissues were analyzed separately only ovaries showed a consistent decline in BODIPY fluorescence with age (Supplementary Table S8, Supplementary Fig. S7). There was also a significant clone-by-age interaction, suggesting that clones differ in the age-related dynamics of lipid hydroperoxides in tissues (Supplementary Fig. S6 E, Supplementary Table S7). However, the slope of such dynamics showed no correlation with clones’ longevity (Supplementary Fig. S6 F). Whole body extract MDA content slightly increased over 3 age classes studied, again, the clone-by-age interaction dwarfing the age effect (Supplementary Fig. S6 G, H).

While we saw little evidence for the overall correlation between LPO parameters and longevity, such correlation appeared to be different among the long-lived, CR-responsive clones and the short-lived CR non-responsive ones (Fig. 5 C – F). There only case in which there was an overall negative correlation with longevity (MDA abundance at the age of 25 days) the same correlation was actually positive within the long-lived clones (Fig. 5E). The latter correlation switched signs in 55 days old *Daphnia* (Fig. 5 F).

## Discussion

An extensive set of lifespan experiments conducted under a variety of laboratory protocols revealed a significant variation in *Daphnia* baseline lifespan caused by both genetic variability and variation of experimental treatments, mimicking similar results previously obtained for nematodes (Lucanic et al. 2017; Urban et al. 2021), *Drosophila* (Ziehm et al. 2013), and mice (Bartke et al. 2019; Spiridonova et al. 2024). Important for the use of *Daphnia* as a model for longevity and aging studies, conspecific isolates (maintained as parthenogenetically reproducing clones) showed a nearly three-fold difference in longevity, similar to the differences observed in interspecific comparisons in *Daphnia* (Dudycha 2003). Consistent with data on inbred depression in small rockpool populations (Lohr et al. 2014; Lohr and Haag 2015), northern habitat, small pond, and rockpool-originated clones had the shortest lifespan, although we were unable to untangle the effect of latitude and habitat type due to a small number of clones originating from each habitat type.

Among laboratory conditions that are likely to vary in lifespan experiments, apart from the obvious effect of temperature, the size of groups in which *Daphnia* are housed emerged as a factor to be aware of, with individually housed *Daphnia* showing the longest lifespan. Non-random clustering of death in experiments with group housing indicates that shortened lifespan under this type of husbandry may be caused by a short-lived infection or a volatile toxin emitting from a carcass. Group size disproportionally affected early survival and had little effect of late-life survival (although this may be caused by cohorts attrition naturally resulting in small group sizes). There was little evidence of clone-by-group size interaction. The use of meshed inserts had no statistically significant effect on longevity across multiple comparisons, although more frequently increased than decreased lifespan in several combinations of group size and clones used (cf. Cho et al 2023 for the use of fully automated neonate removal using inserts). Medium composition, at least the two commonly used reconstructed pond water formulations tested, did not radically affect the mean lifespan over all clones tested, affected late-life survival stronger than early-life survival and showed a significant clone-by-water type interaction. If anything, the higher hardness ADaM medium results in slightly longer lifespan than the lower Ca^2+^ containing COMBO, which may be the reason why some researches opt for a high-hardness modification of COMBO (Constantinou et al. 2019).

An effect similar in magnitude and presence of strong interactions was demonstrated by the two species of food algae – a green alga *Scenedesmus* and a stramenopile alga *Nannochloropsis,* which is, in contrast to green algae, known to be rich in PUFAs. It remains unknown if the use of other, less commonly used algal species, such as *Chlamydomonas reinhardtii* (Pietrzak et al. 2018) or *Chlorella vulgaris* (Constantinou et al. 2019) would produce a lifespan altering effect in a common-garden experiment. An attempt to replace freshly grown algal food by commercially available dry cyanobacterial diel, which could have been advantageous in term of diet standardization, demonstrated that this is probably going to be difficult to achieve without a significant decrease of longevity, i.e., without possible masking of beneficial longevity-extending intervention effects.

Interestingly, the lifespan differences between identical experiments conducted by different investigators simultaneously and in the same lab may be greater than those between similar experiments conducted by the same investigator in different labs.

One additional variation of the lifespan experiments that deserves consideration is the sex of experimental animals. Using male-only cohorts eliminates the need of neonate removal (and of course any possibility of Jeanne Calment effect), but also takes any possible trade-offs with the large energy expenditure of producing eggs out of the consideration and reduces generality of finding if the two sexes age differently, and respond to interventions differently, i.e., there are sex-by-age or sex-by-treatment interactions. Males in *Daphnia* consistently show a shorter lifespan than females (Constantinou et al. 2019; but see Pietrzak et al. 2018), an observation made nearly a hundred years ago (Macarthur and Baillie 1926) that served as one of the first evidence supporting the rate of living hypothesis (Rubner 1908; Pearl 1928; Speakman 2005). Considering the more recent interpretation of the rate of living hypothesis in terms of the amount of oxidative damage accumulated over the lifetime as the limiting factor for lifespan extension (Austad 2018), one would expect the shorter lifespan in *Daphnia* males to be associated with faster, in astronomical time, but similar, over relative age, accumulation of reactive oxygen species-mediated damages. Indeed, the amount of DNA damage followed this pattern in *Daphnia* males and females (Constantinou et al. 2019), but the accumulation of the final product of lipid peroxidation, MDA, did not: it appears to accumulate in males slower, not faster, than in females (Coggins et al. 2017; Constantinou et al. 2019), an effect possibly related to lipid composition differences between sexes (Constantinou et al. 2020; see below). No combination of our 25 experiments provided an opportunity to test for clone-by-sex interactions or for any other interaction between sex and protocol parameters except group size (which was not significant). Thus it remains to be tested if males react to experimental treatments and longevity interventions differently from females. There is evidence that such differences may be clone- (Pietrzak et al. 2018) and temperature-dependent (Pietrzak et al. 2018; this study).

Considering a significant reduction of lifespan in food levels above saturation observed in this study and many others, shorter lifespan in males can very well be a manifestation of the same effect. Adult females 5-8 times larger than adult males by biomass, and males also have lower reproductive expenditures, so maintaining both sexes at the same food level (Pietrzak et al. 2018; Constantinou et al. 2019) may actually constitute a CR experiment, with females experiencing below-saturation food availability, while males – above saturation one. In most male cohorts analyzed here (except for the caloric restriction experiment) males have been maintained at 3x density relative to females, but this may still not be enough to compensate for biomass and expenditure differences.

Consistent with the classic dietary restriction work (Ingle 1933; Ingles et al. 1937; Lynch and Ennis 1983) we show a clear caloric restriction effect in *Daphnia*. However, pronounced life-shortening effects of *ad libitum* feeding start to be observed only at fairly high food concentration, at least for the large body size species like *D. magna.* This clarifies the reasons why more recent work (Schwartz et al. 2016) failed to observe this effect. Although Schwartz et al. (2016) do no report absolute food concentrations used, only dilutions of a maximal level (an example of experimental conditions under-reporting that makes longevity studies hard to compare) one can conclude, from fecundity values reported, that even the highest food level used was not truly *ad libitum*. Furthermore, we show a significant genotype-by-environment interaction in the caloric restriction effect (providing a further reason why some studies might not be able to observe it), with short-lived clones showing no life extension in dietary restricted conditions. This may be an intra-specific continuation of the same pattern observed in the between-species comparison (Dudycha et al. 2003). However, the caloric-restriction differences between the short-lived *D.pulex* and long-lived *D.pulicaria* are likely to be the result of different selection regimes in different habitats (Dudycha et al. 2003), the interspecific differences observed here appear to be a manifestation of inbreeding in small intermittent (i.e., frequently re-established from a small number of founders) populations (Lohr et al. 2014; Lohr and Haag 2015): most of our short-lived, non caloric restriction-responsive clones originated from small intermittent habitats. A recent study that also reported that the caloric restriction life extension is not observed in genotypes genetically predetermined for short life has also recently been observed in *Drosophila* (Sember et al. 2024) interprets this as deficiency in DR-detecting signaling pathways. However, there can be a plethora of other reasons for this observation, including deficiencies in antioxidant mechanisms triggered by DR, as suggested by our LPO results on the differences in LPO between the long-lived, caloric restriction-responsive and short-lived, non-caloric restriction-responsive clones.

Despite the lack of support for the oxidative damage as the explanation of shorter lifespan in males, lipid peroxidation in *Daphnia* has emerged as a hallmark of aging (Barata et al. 2005) and a strong correlate of aging rate in among-clone comparisons (Ukhueduan et al. 2022). There are two possible causes of lipid peroxidation level change across ages, diets, or genotypes: the amount of ROS and the amount of their lipid targets, namely PUFAs, both of which can change with age. It appears that lipid composition changes significantly with age (Barata et al. 2005), markedly differently in males and females (Constantinou et al. 2020), with, in particular, highly polyunsaturated fatty acids declining with age (Barata et al. 2005). Furthermore, the amount of LPO and its age trajectory may differ within each ovary cycle as lipids, including PUFAs are accumulated in tissues and transported into the ovaries (Lowman and Yampolsky 2023; Campbell and Yampolsky, in preparation). In this study we tried to untangle the standing crop of lipid hydroperoxides from the accumulation of the final product of LPO by measuring these two parameters, respectively, by in-situ fluorescence microscopy of tissues stained in-vivo with BODIPY C11 marker of lipid hydroperoxide and by whole body extract TBARS determination of MDA content. Both these parameters are affected by two quantities: the abundance of oxidizing agents (e.g. ROS) and the abundance of oxidation targets (among lipids primarily PUFAs). However, they measure two different features of lipid peroxidation biochemistry.

BODIPY fluorescence (specifically the ratio of green fluorescence to the sum of green and red fluorescence) measures the relative abundance of peroxidized lipids (normalized by total lipid content in the tissue) at the time of measurement. MDA content, on the other hand, shows total amount of final product of lipid peroxidation, which can accumulate over time. In this study, both parameters showed statistically significant, but slight relationships with age which are of the opposite directions: BODIPY fluorescence decreased with age, while MDA increased. Both effects were dwarfed by the abundance of clone-by-age interactions. For both effects, these interactions notwithstanding, there was no correlation between the rate of change over age and clone-specific longevity. Neither parameter appeared to correlate with lifespan in short-lived clones and both showed a positive correlation with lifespan among the long-lived clones, a relationship that was reversed for the MDA concentration later in life. We interpret this as an evidence of beneficial effects of PUFAs accumulation in tissues early in life, which is offset by the accumulation of toxic product of LPO later in life. In other words, the most long-lived clones tend to accumulate PUFAs in their tissues, but to be capable to avoid accumulation of MDA.

Given the clonal and laboratory condition lifespan differences described above, in order to generate comparable results suitable to provide a control longevity baseline for intervention studies, *Daphnia* longevity experiments should be conducted in standardized conditions, and on a standard reference genotype, or, better, several genotypes. We propose that if maximal possible lifespan is desired in the control cohort such standards should include the use of ADaM medium, *Scenedesmus* algae at food, 20-25 °C temperatures. Ideally individual housing of experimental animals should be used, or, when practical reasons demand *Daphnia* to be maintained in groups, the use of meshed inserts to eliminate manual handling of individuals and neonate removal should be encouraged. Ideal clones for this purpose are those that show high longevity under a variety conditions and a stronger caloric restriction effect on longevity if these effects are relevant.

## Data and material availability

Supplementary data (REF) include a summary table with median and mean lifespan values observed in all experiments in all clones and raw survival data for each individual experiment. All *Daphnia* clones listed in Supplementary Table S1 are available both from Basel University *Daphnia* stocks collection or from LYY lab, upon request.

## Acknowledgments

We are grateful to authors of previously published experiments included into this study, including Cora E. Anderson, Bret L. Coggins, Millicent N. Ekwudo, Taraysha D. Moore, and Yongmin Cho; to Dieter Ebert and Jürgen Hottinger for supplying *Daphnia* clones, and to Mark W. Kirschner for stimulating discussions during early stages of this work. This work was supported by Impetus Foundation grant to LYY and several ETSU Student-Faculty Collaboration grants and Small RDC grants.

