## Supplementary Figures and Tables for "Short lifespan is one’s fate, long lifespan is one’s achievement: lessons from *Daphnia*"

**Summary of experiments and clones**

Table S1. List of lifespan experiments included in the analysis. ExpID: unique identifier of an experiment (see Supplementary materials) Locations: “HMS” – Kirschner lab at Harvard Medical School; “ETSU” – Yampolsky lab at East Tennessee State University. Nc: Number of independent cohorts. V: container volume. Food species: *S.a. - Scenedesmus acutus^1^; N.l. - Nannochloropsis limnetica*. C_f_: Food concentrations used, cells/mL/day. References listed if previously published.

| ExpID | Location | Nc | Group size | V, mL | Container type, Neonate removal method | Inserts used? | Insert volume | Water | T °C | Clones used | Male cohorts | Food species | C_f_ | Other treatments in the same experiment ^8^ | Ref. |
| --- | --- | --- | --- | --- | --- | --- | --- | --- | --- | --- | --- | --- | --- | --- | --- |
| 1 | HMS | 6 | 5 | 100 | jars, manual | N |  | COMBO | 25 | IL | Y | *S. a.* | 50 / 100 |  |  |
| 2 | ETSU | 1 | 5 | 100 | jars, manual | N |  | COMBO | 25 | several | Y | *S. a.* | 100 |  |  |
| 3 | ETSU | 1 | 10 | 200 | jars, semi-manual | Y | 100 | COMBO | 20 | FI,GB,IL,HU | N | *S. a.* | 100 | DNP exposure |  |
| 4 | HMS | 3 | 50 | 1000 | SmartTanks, auto | Y | 1000 | ADaM | 25 | IL | N | *S. a.* | 100 | various | 2 |
| 5 | ETSU | 2 | 20 | 400 | tanks w. 8-cup inserts, semi-manual | Y | 100 | COMBO | 20 | FI,GB,IL,HU | N | *S. a.* | 100 | intermittent hypoxia | 3 |
| 6 | ETSU | 1 | 5 | 100 | jars, manual | N |  | COMBO | 20 | FI,GB,IL,HU | Y | *S. a.* | 25 / 100 | L/H food switch |  |
| 7 | ETSU | 3 | 5 | 100 | jars, manual | N |  | COMBO | 20 | FI,GB,IL,HU | N | *S. a.* | 100 |  | 4 |
| 8 | ETSU | 1 | 1 | 20 | vials, manual | N |  | COMBO | 20 | FI,GB,IL,HU | N | *S. a.* | 100 | maternal exposure to hypoxia | 5 |
| 9 | ETSU | 2 | 5 | 100 | jars, manual | N |  | COMBO | 20 | FI,GB,IL,HU | N | *S. a.* | 100 | old maternal age | 6 |
| 10 | HMS & ETSU | 1 | 1 | 20 | vials, manual | N |  | COMBO | 20 | FI,GB,IL,HU | N | *S. a.* | 100 | old maternal age | 6 |
| 11 | ETSU | 1 | 50 | 1000 | tanks, semi-manual | Y | 1000 | ADaM | 20 | GB,IL | N | *S. a.* | 100 / 200 | exposure to BHB,NMN |  |
| 12 | ETSU | 1 | 1 | 20 | vials, manual |  |  | ADaM | 20 | FI,GB,IL | N | *S. a.* | 100 | Maternal BHB exposure |  |
| 13 | ETSU | 1 | 10 | 200 | jars, semi-manual | Y | 100 | ADaM | 20 | GB, IL | Y | *S. a.* | 25 / 50 / 100 / 200 | exposure to BHB |  |
| 14 | ETSU | 1 | 10 | 200 | jars, semi-manual | Y | 50 | ADaM | 20 | FI,GB,IL,HU | N | *S. a.* | 100 |  |  |
| 15 | ETSU | 9 | 5 | 100 | jars, manual | N |  | ADaM | 20 | IL | Y | *S. a.* | 100 |  |  |
| 16 | ETSU | 6 | 10 | 200 | jars, semi-manual | Y | 50 | ADaM | 20 | FI,GB,IL | Y | *S. a.* | 100 |  |  |
| 17 | ETSU | 1 | 1 | 20 | vials, manual | N |  | COMBO, ADaM | 20, 24 | FI,GB,IL,HU | N | *S. a.* | 100 |  |  |
| 18 | ETSU | 1 | 5 | 100 | jars, semi-manual | Y^10^ | 50 | ADaM | 20 | 12 clones | N | *S. a.* | 100 |  |  |
| 19 | ETSU | 1 | 5 | 100 | jars, semi-manual | Y | 50 | ADaM | 20 | GB, IL | N | *S. a.* | 50 / 100 / 200 / 300 / 400 |  |  |
| 20 | ETSU | 1 | 5 | 100 | jars, semi-manual | Y | 50 | ADaM | 20 | 12 clones | N | *S. a.* | 50 /100 / 400 |  |  |
| 21 | ETSU | 1 | 1 | 20 | vials, manual | N |  | ADaM | 12, 20, 28 | FI,GB,IL,HU | N | *N.l.* | 100 | Food growth T |  |
| 22 | HMS | 1 | 1 | 100 | jars, semi-manual | Y | 10 | ADaM | 21 | IL | Y | *S.a.* | 100 | Dry Spirulina as food |  |
| 23 | ETSU | 1 | 10 | 200 | jars, semi-manual | Y | 100 | COMBO | 20 | GB, IL | N | *S. a.* | 12.5 / 25 / 50  / 100 / 200 |  |  |
| 24 | ETSU | 1 | 5 | 100 | jars, manual | N |  | COMBO | 20 | FI,GB,IL,HU | N | *S. a., N.l.* | 20 / 200 |  | 7 |
| 25 | ETSU | 1 | 1 | 200 | vials, manual |  |  | COMBO | 25 | Several^11^ | N | *S. a., N.l.* | 100 |  | 9 |

Notes and references:

1 Correct current taxonomy *Tetradesmus obliquus*

8. Not used in the current analysis.

9. Coggins B.L., A.C. Pearson, L.Y. Yampolsky. 2021. Does geographic variation in thermal tolerance in Daphnia represent trade-offs or conditional neutrality? Journal of Thermal Biology 98: 102934. https://doi.org/10.1016/j.jtherbio.2021.102934

10. Neonates removed every other day, not simultaneously with water change as in other experiments.

11. Too few replicates per clone, only combined data analyzed.

Table S2. Provenance of clones used in the experiments included into the analysis. 2-letter code of the 4 reference clones used elsewhere in the text or figures it is shown in bold here. Source of data: Dieter Ebert, personal communication, amended.

| Clone ID  (abbreviated ID) | Latitude | Longitude | Site description | Season with plankton | Summer dry | Winter freezing | Longest dimension, m | Max depth, cm | Elevation, m | Rock-pool | Estuary/ coastal | Salinity category | Sampled by |
| --- | --- | --- | --- | --- | --- | --- | --- | --- | --- | --- | --- | --- | --- |
| DE-KA-F28  (DE) | 50.935361 | 6.927944 | Aachener Weiher city pond, Köln, Germany | summer | no | no | 152 | 160 | 52.4 | 0 | no | fresh | Eric von Elert |
| DK-RL-3  (DK) | 55.964167 | 9.596389 | Ring Lake, Braedstrup, Denmark | year around | no | yes | 924 |  | 78 | 0 | no | fresh | Luc DeMeester |
| FI-FAV-1-1  (FI-FAV) | 60.021612 | 19.902506 | Aland Island/ Vestra Masskaer rockpool, Finland | summer | yes | yes | 5 | 30 | 0.8 | 1 | coastal | rockpool | Thomas Zumbrunn |
| **FI**-FSP1-16-2  (FI-FSP) | 60.184363 | 25.794781 | Suur Pellinki Skerry islands, Finland | summer | yes | yes | 2 | 30 | 1 | 1 | coastal | rockpool | Dieter Ebert |
| FR-TR-1  (FR) | 43.738306 | 3.863302 | Pond, Le Triadou, France | winter | no | no | 70 |  | 74.4 | 0 | no | probably fresh | Christoph Haag |
| **GB**-EL75-69  (GB) | 51.527556 | -0.158147 | Regents Park Lake, London, UK | year around | no | no | 914 |  | 31.8 | 0 | no | fresh | Daniela Brunner |
| **HU**-K-6  (HU) | 46.796389 | 19.185 | Kiskunsagi National Park lake, Kelemen-szek, Hungary | summer | no | yes | 1524 |  | 89 | 0 | no | salty | Luc DeMeester |
| **IL**-M1-8  (IL) | 31.778213 | 35.220601 | Mamilla Pond, Jerusalem, Israel | winter | yes | no | 96 |  | 777.9 | 0 | no | fresh | Frida Ben-Ami |
| MA-ES-3  (MA) | 31.490714 | -9.76443 | Small pond near Essaouira, Marocco | winter | yes | no | 10 |  | 1.8 | 0 | Coastal marsh | coastal marsh | Nadja Brun |
| RU-BOL1-1  (RU-B) | 66.4213833 | 33.8485861 | Bol. Astafiy Island near White Sea Biol. Station, Russia | summer | yes | yes | 5 | 30 | 3 | 1 | coastal | rockpool | Sergey Glagolev |
| RU-PIK-2  (RU-P) | 59.994906 | 30.473075 | Pond, St. Petersburg, Russia | summer | no | yes | 457 |  | 20.3 | 0 | no | fresh | Yan Galimov |
| RU-VOL-36  (RU-V) | 48.53 | 44.486944 | Pond, Volgograd, Russia | spring & autumn | yes | yes | 200 | 200 | 5 | 0 | no | salty | Yan Galimov |

**Median lifespan correlates: geography of clones’ origin**

A	B

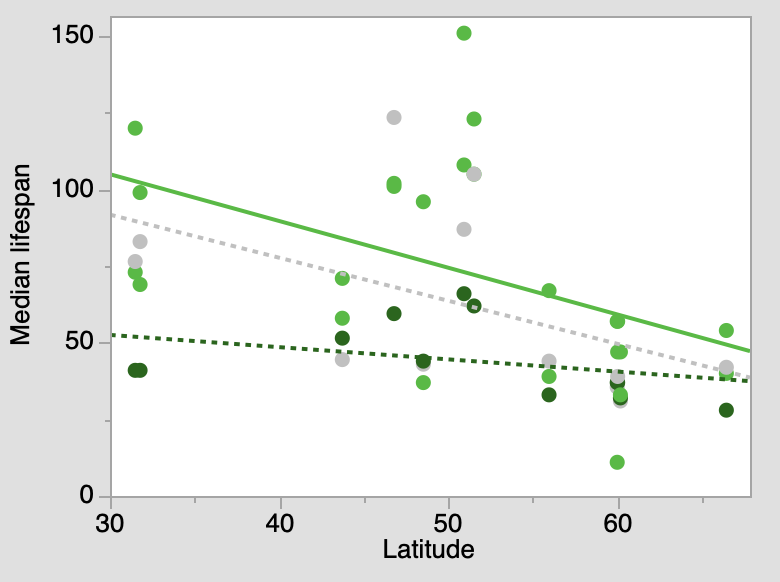

P < 0.02

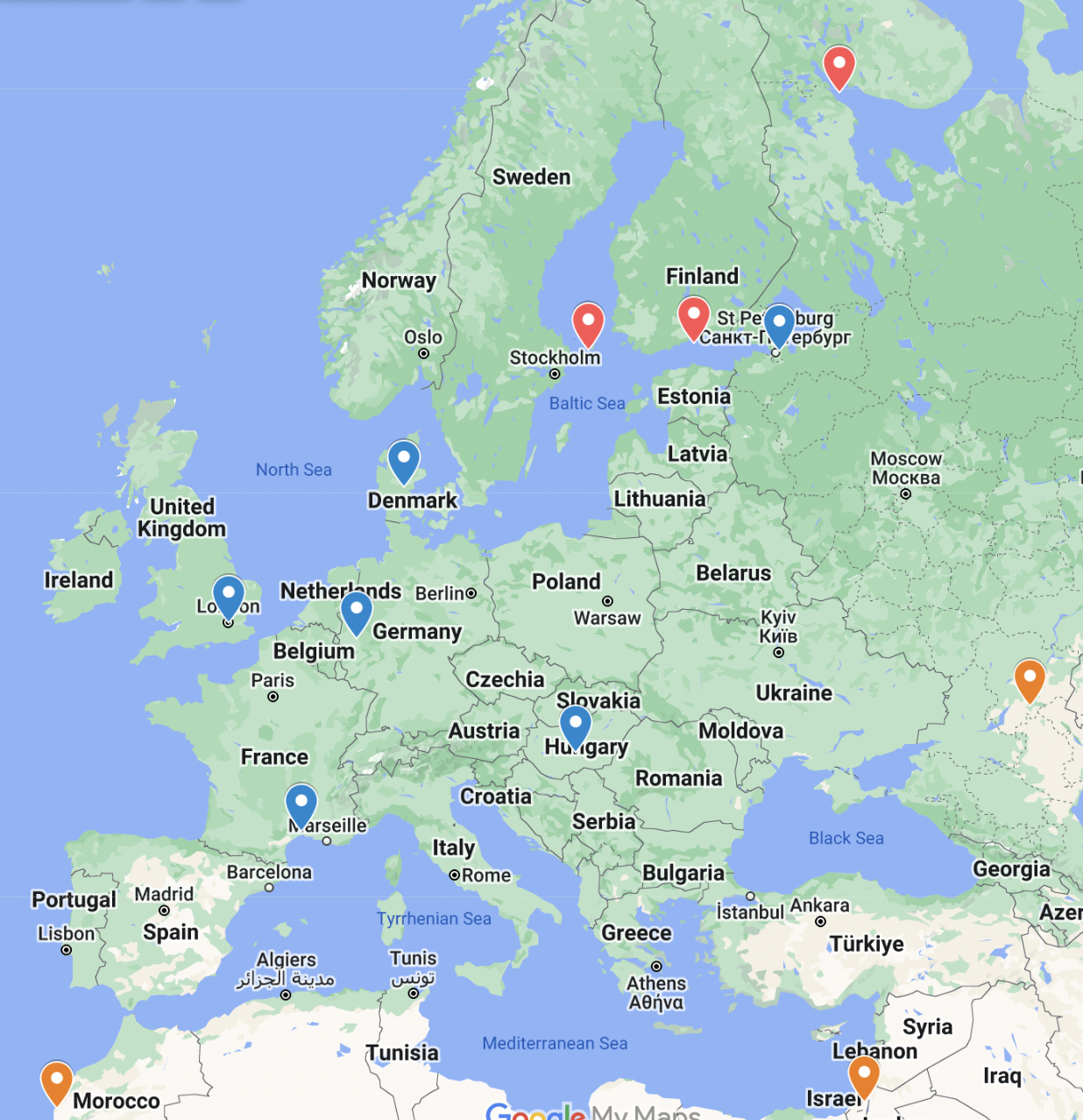

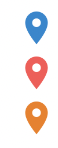

Permanent pond or lake

Intermittent rock pool

Intermittent summer dry pond

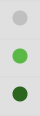

50

100

400

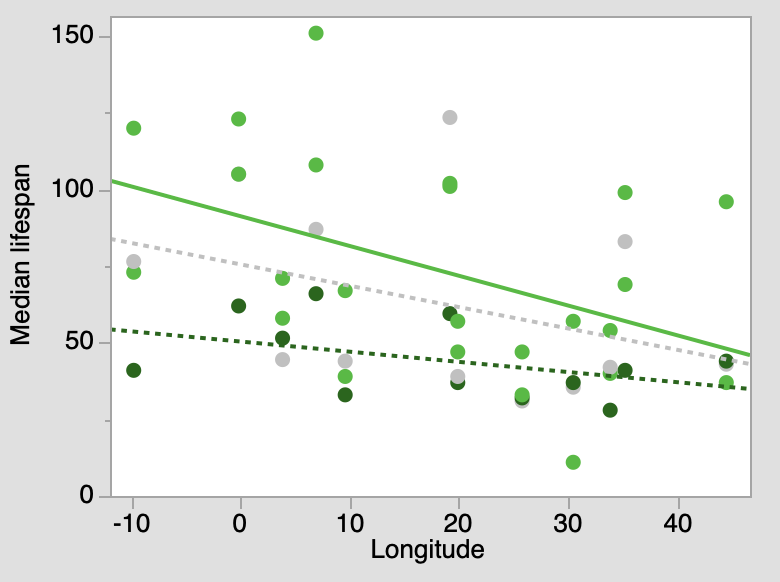

P < 0.03

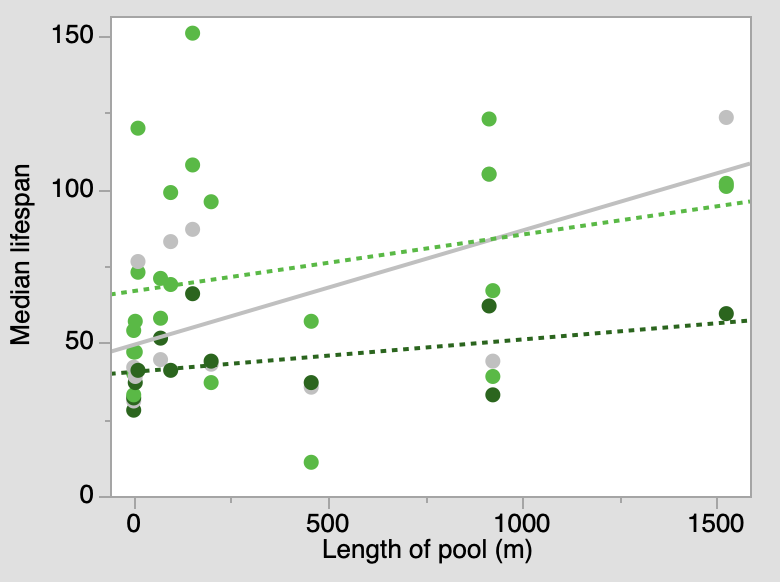

P < 0.04

C

D

E	F	G

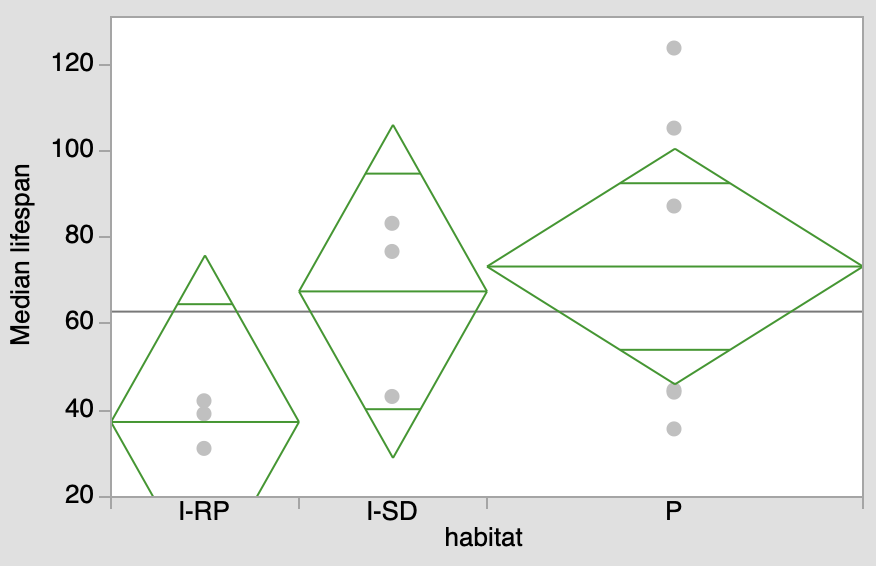

P > 0.2

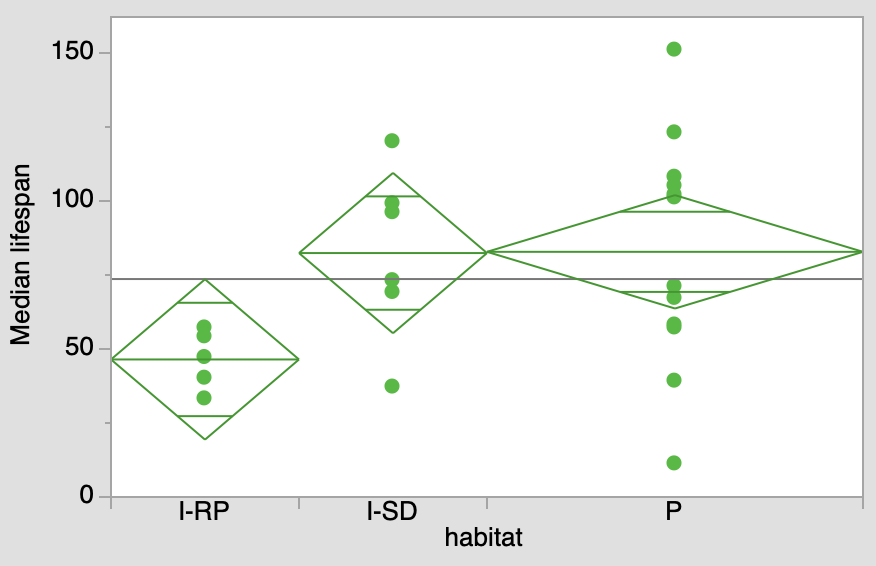

P > 0.075

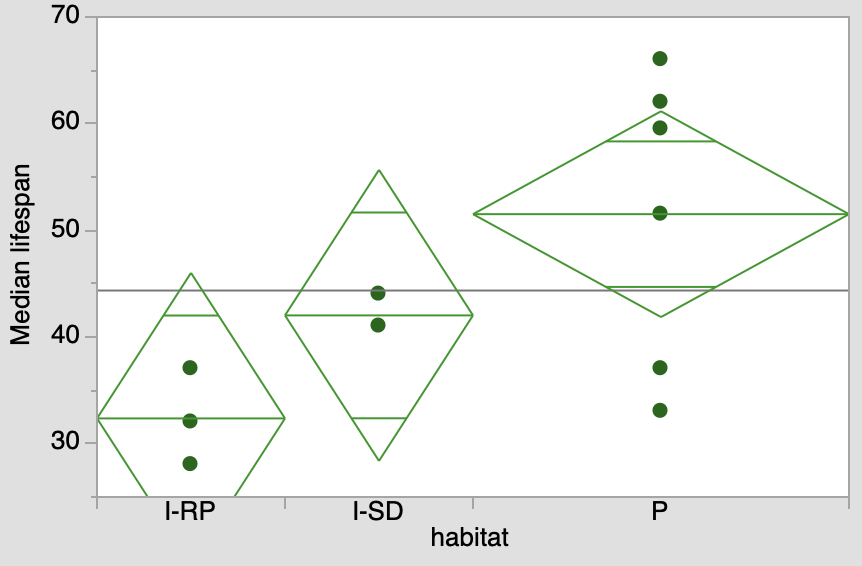

P > 0.075

Fig S1. Geographical origins of 12 *D. magna* clones (A) and geographic correlates of clones’ median lifespan measured at 3 food levels (50, 100, and 400 10^3^ cells/mL/day). B, C: correlations with latitude and longitude; D: correlation with habitat size (longest dimension); E – G: lack of significant differences between habitat types (I-RP, intermittent rockpool; I-SP, intermittent, summer dry; P, permanent lake or pond).

Fig. 1. Geographic origin of clones used in the experiments (left panel) and clones’ survival curves (right panel)

**Median lifespan correlates: group size, sex, food level.**

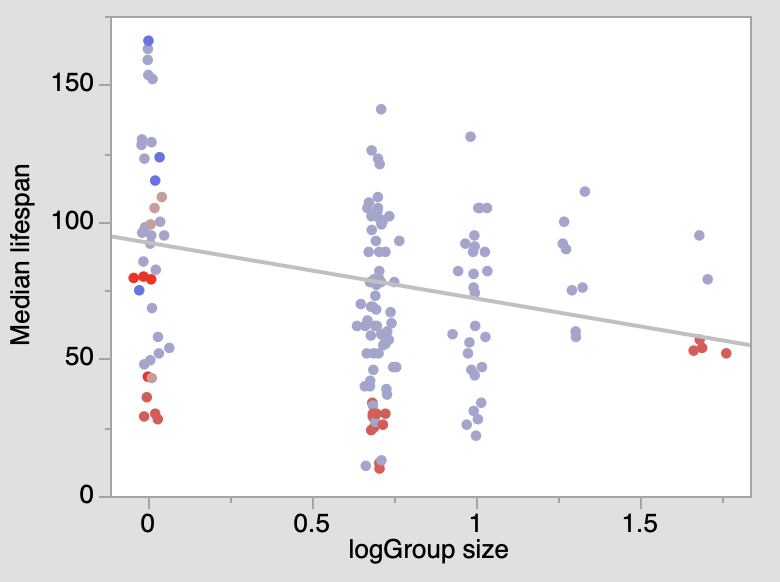
A	B

Median lifespan

|  | No inserts | Inserts |
| --- | --- | --- |
| FI 5 in 100 | 51.1429 | 44 |
| GB 5 in 100 | 71 | 102.333 |
| HU 5 in 100 | 71.2143 | 100.5 |
| IL 5 in 100 | 64.9524 | 71 |
| GB 10 in 200 | 51.5 | 84.8333 |
| IL 10 in 200 | 57 | 77.1 |

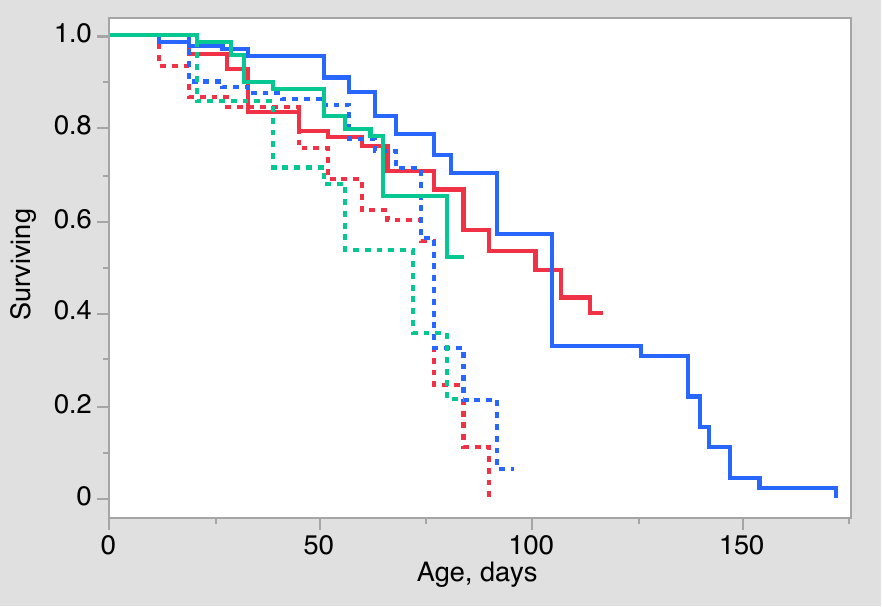
C	D

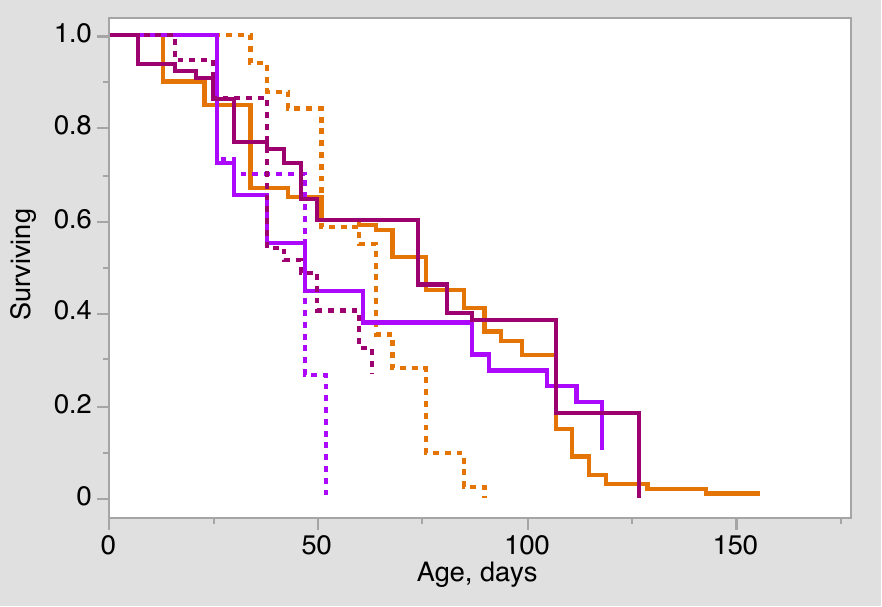
E

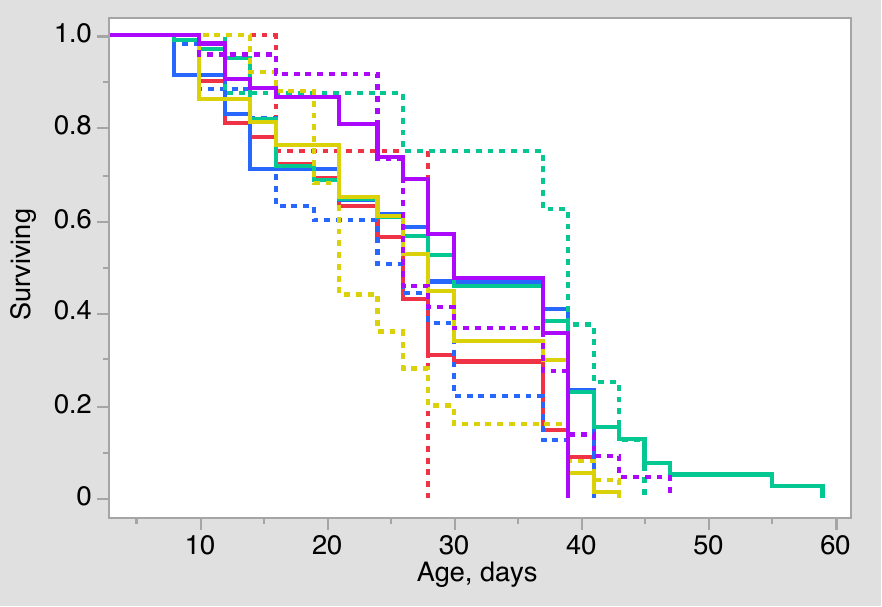

| Source | DF | L-R c2 | P |
| --- | --- | --- | --- |
| Clone | 4 | 9.54 | 0.05 |
| sex | 1 | 0.40 | 0.53 |
| Clone*sex | 4 | 2.05 | 0.73 |

Fig. S2. A: as main text Fig.1A, but all clones included. B: Effect of the use of inserts (semi-manual neonate removal) on median lifespan in 4 reference clones under 5 individuals in 100 mL setup and in 2 clones under 10 individuals in 200 mL setup. No comparison survives Tukey test (P > 0.05). C, D: as main text Fig.1 B, but independent cohorts within each experiment analyzed separately; solid lines: females, dotted lines: males. E: As main text Fig. 2, females (solid lines) vs. males (dotted lines) in an experiment conducted at 25^o^C. Proportional Hazards analysis failed to converge, Weibull (best) distribution parametric survival analysis results shown.

[[Experiment 2, 25^o^C]]

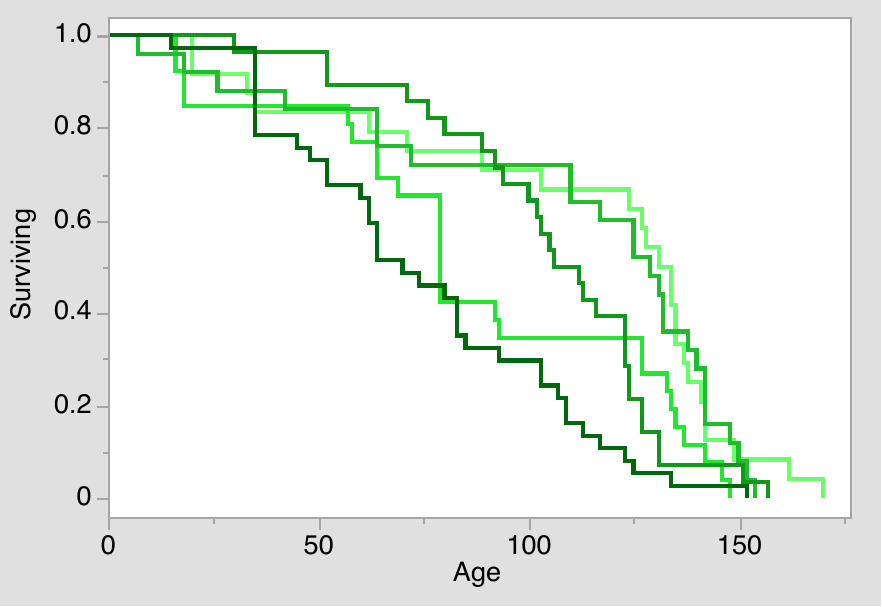

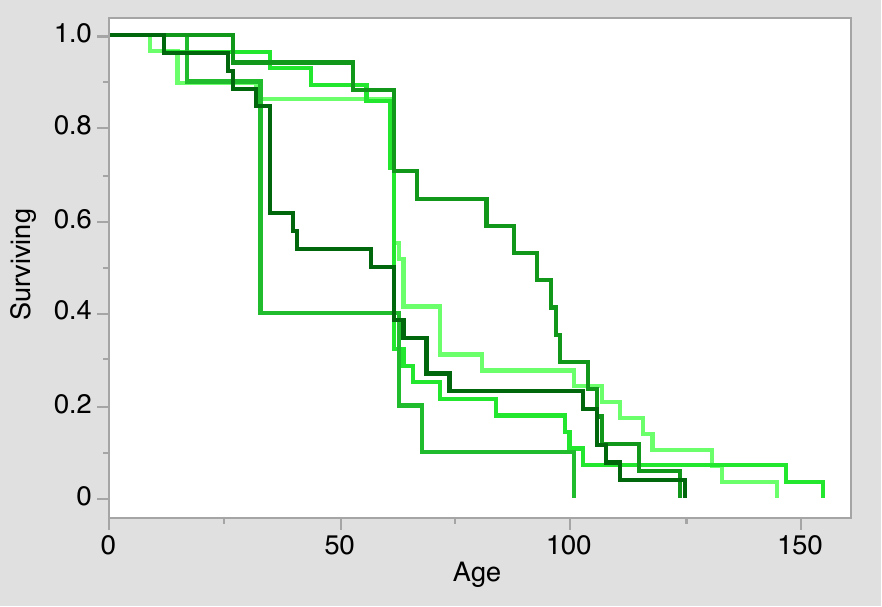

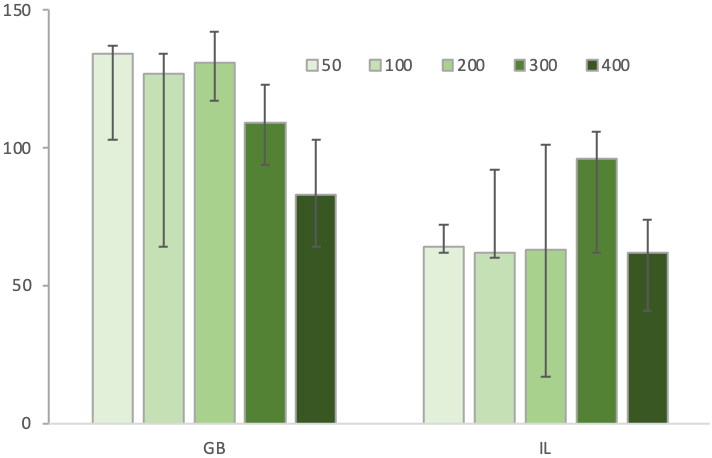

| Source | DF | χ2 | P |
| --- | --- | --- | --- |
| Food | 4 | 12.7 | 0.015 |
| Clone | 1 | 17.2 | 0.0001 |
| Food*Clone | 4 | 5.6 | 0.23 |

|  | 50 | 100 | 200 | 300 | 400 |
| --- | --- | --- | --- | --- | --- |
| GB | 134 | 127 | 131 | 109 | 83 |
| IL | 64 | 62 | 63 | 96 | 62 |

Fig. S3. Data similar to main text Fig. 4A,B from an experiment with 2 clones maintained at 5 food levels. Food concentration x 10^3^ cells/mL/day. Insert: proportional hazards analysis results.

[[ExpID=19]]

Table S3. Proportional hazard analysis (Log-likelihood test) of the effect of food level (0.125, 0.25, 0.5, 1, and 2 x E5 cells/mL/day) in two sexes in two clones (GB and IL). [[Exp 23]]

| Source | DF | L-R ChiSquare | Prob>ChiSq |
| --- | --- | --- | --- |
| food level | 4 | 66.35 | <0.0001 |
| sex | 1 | 5.06 | 0.024 |
| food level*sex | 4 | 42.34 | <0.0001 |
| clone | 1 | 0.3 | 0.58 |
| food level*clone | 4 | 5.49 | 0.24 |
| sex*clone | 1 | 5.26 | 0.02 |
| food level*sex*clone | 4 | 29.26 | <0.0001 |

**Age-related changes in 12 *D. magna* clones: fecundity, feeding rate, and lipid peroxidation**

Fecundity was measured by removing and counting neonates every other day from jar containing 5 *Daphnia* females (at the start of the longevity experiment) or 1 – 5 females as cohort attrition accumulated. The number of ephippia produced in the same time period also recorded. Subitaneous (parthenogenetic) fecundity was then calculated as the number of live neonates produces divided by the number of females alive during the collection period minus the number of ephippia produced (because each female producing an ephippium in a given ovary cycle does not produce asexual offspring in the same cycle). Because of synchronization of ovary cycles in females of the same age, particularly in young ages, sequential 2-day neonate counting periods show a massive negative autocorrelation. Therefore a 6-day sliding window averaging was applied, with the data for each age and each jar being the unweighted average of the data from this, the previous, and the following 2-day counting period.

Table S4. ANOVA of age-related changes in fecundity (6 days sliding average of the number of subitaneous (asexual) offspring per asexual female per day, in 12 *D.magna* clones, ages 7 – 70 days. Cf. Fig. S3.

| Source | DF | Sum of Squares | F Ratio | Prob > F |
| --- | --- | --- | --- | --- |
| Age | 1 | 89.62 | 196.58 | <0.0001 |
| Clone | 11 | 72.09 | 14.38 | <0.0001 |
| Age*Clone | 11 | 23.92 | 4.77 | <0.0001 |
| Error | 1598 | 728.57 |  |  |

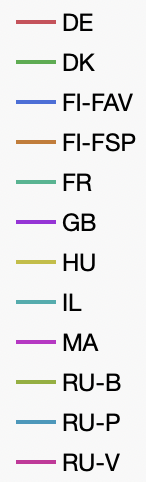

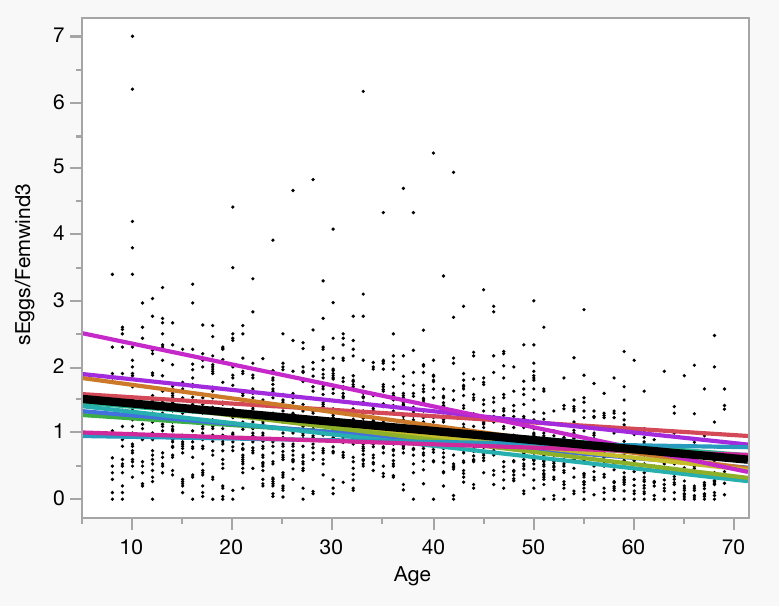

Fig. S4. Age-related changes in fecundity (6 days sliding average of the number of subitaneous (asexual) eggs per asexual female per day, in 12 *D. magna* clones, ages 7 – 70 days. Cf. Table S3.

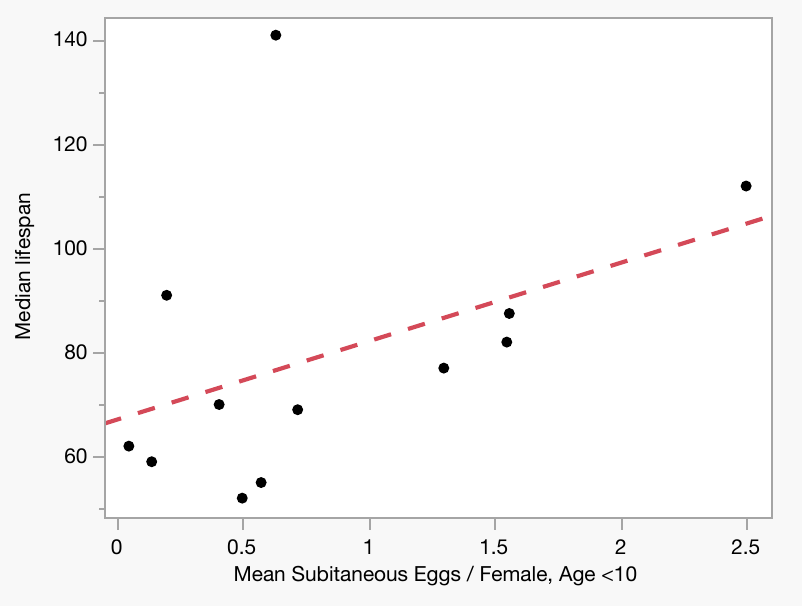

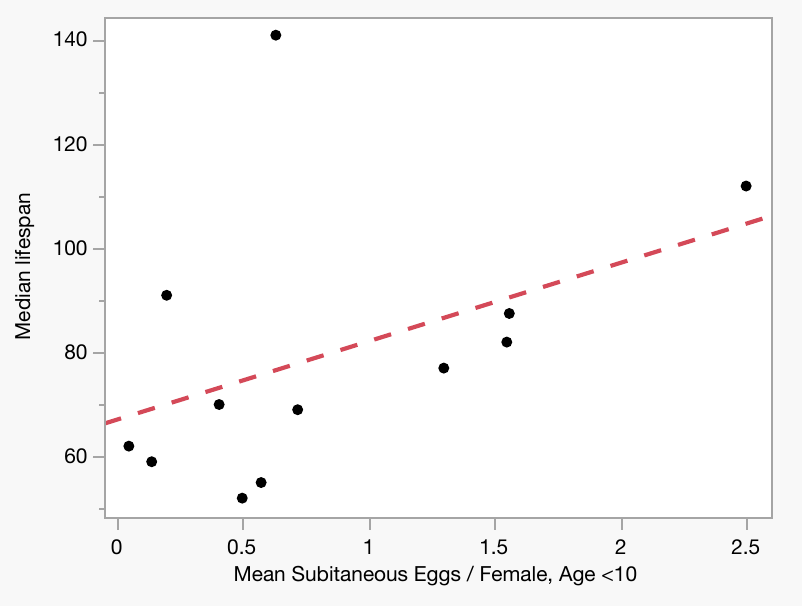

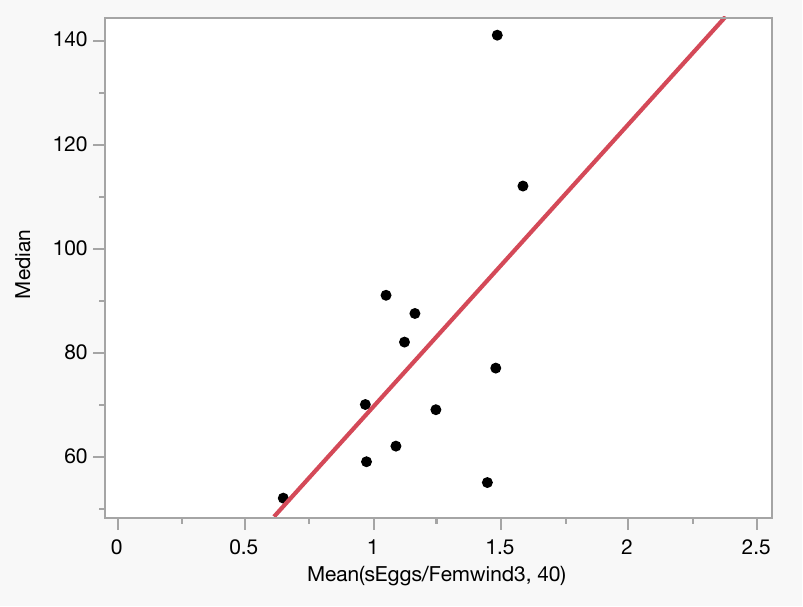

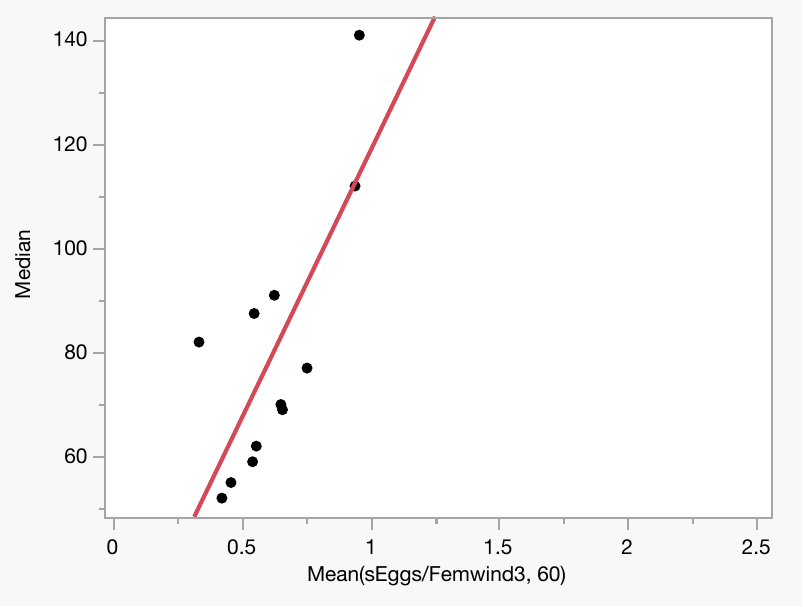

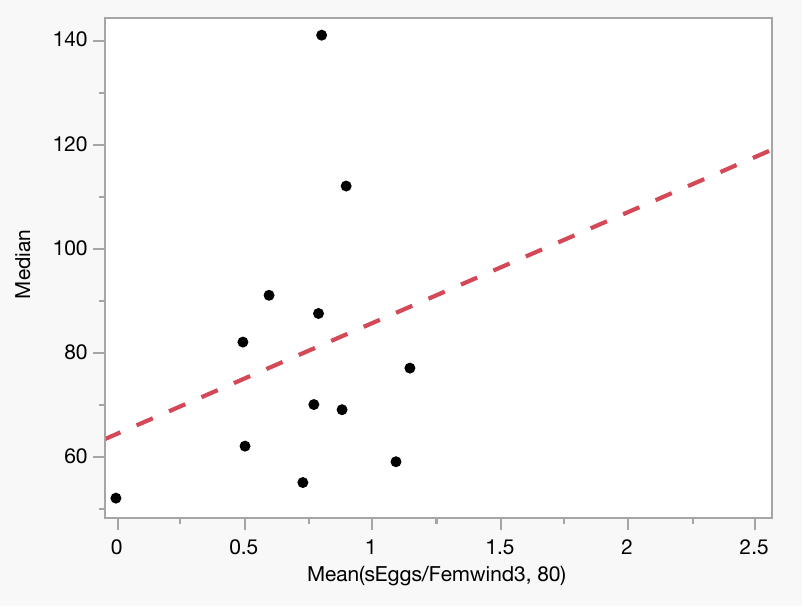

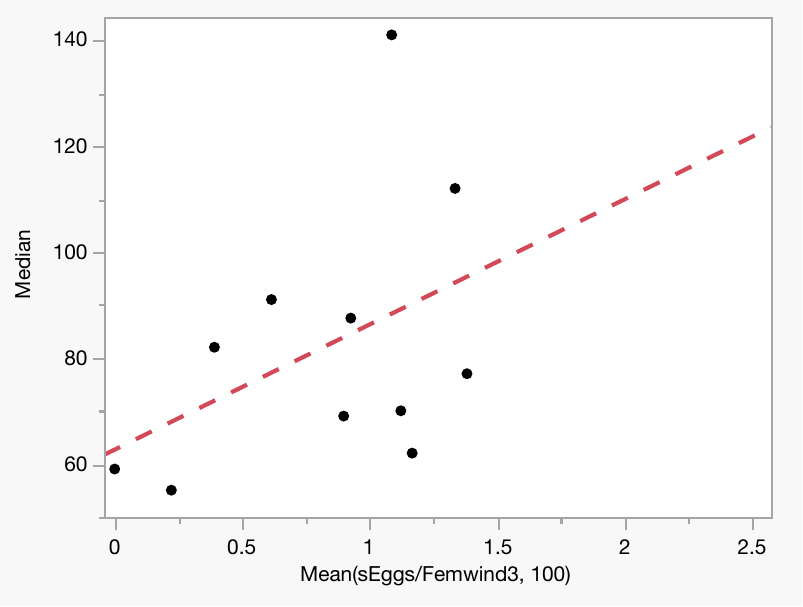

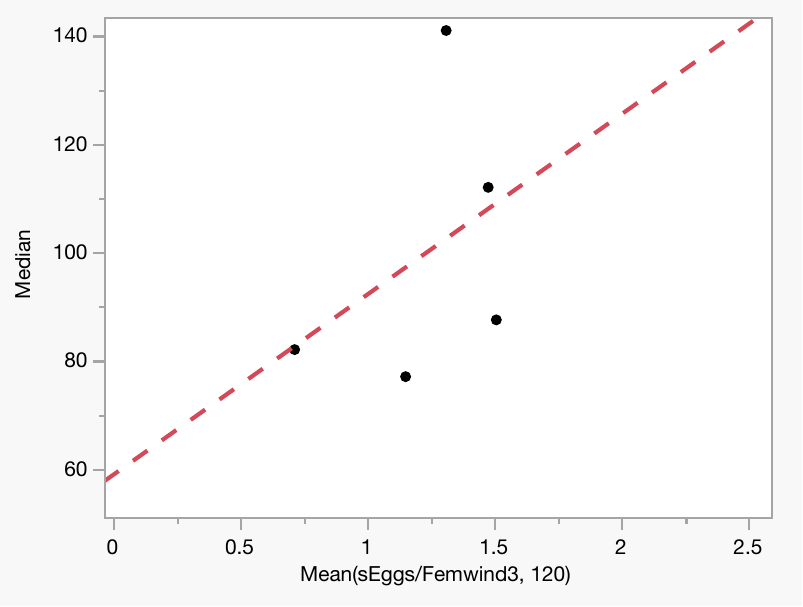

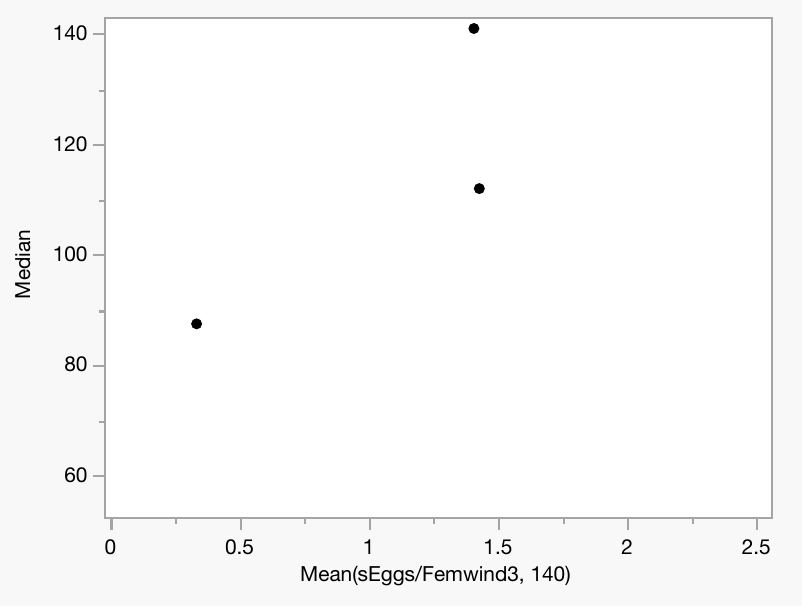

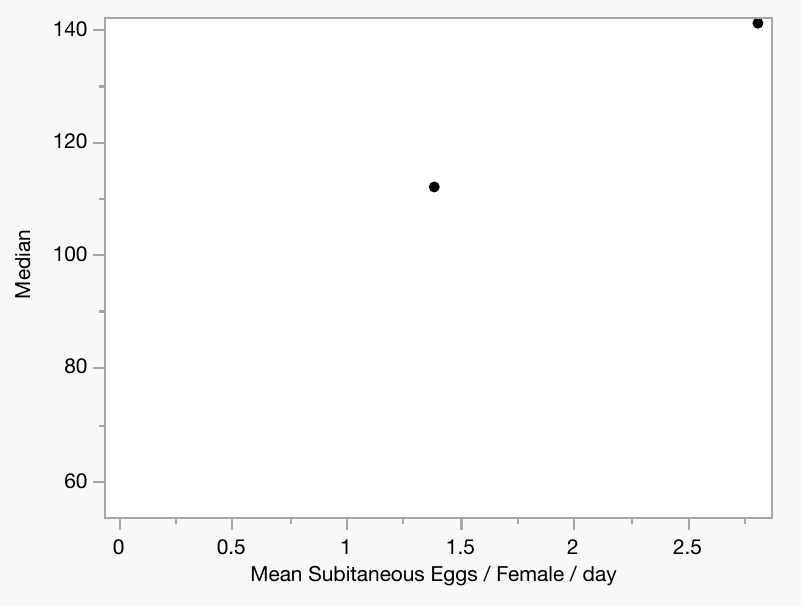

Age < 10

Age 10 - 30

Age 30 – 50

P < 0.05

Age 50 - 70

P < 0.005

Age 70 - 90

Age 90 - 110

Age 110 - 130

Age 130 - 150

Age 150 - 170

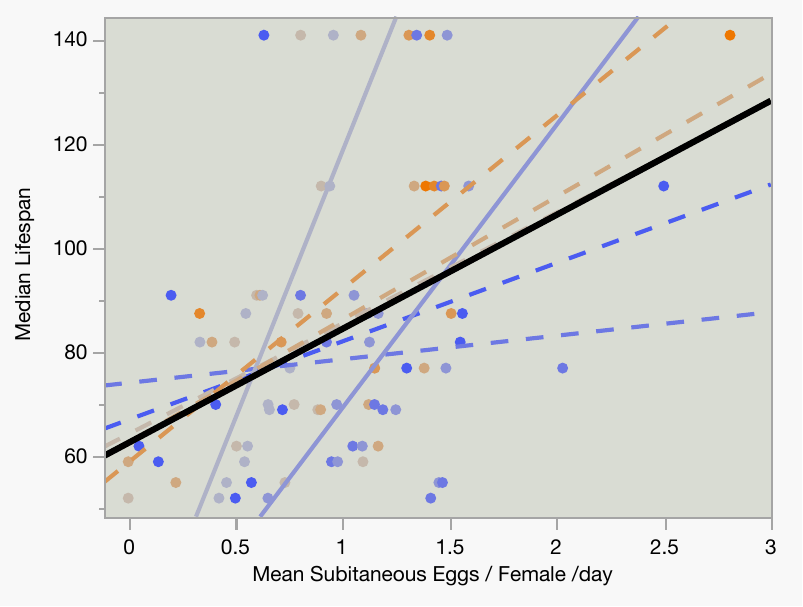

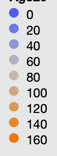

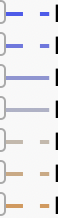

Fig. S5. Lack of fecundity-longevity trade-off in 12 *D. magna* clones. Left: for each of 20-day age classes separately. Right: combined data colored by age (blue to orange). Fecundity: 6 days sliding average of the number of subitaneous (asexual) offspring per female per day. Longevity: median lifespan. Regression shown for age classes to which > 3 clones survive. Regressions that are not significant: dashed lines. Solid black line: regression across all age classes

Table S5. Feeding rate age and clones effects, measured per individual and per mg wet weight. Top: all ages, all clones, bottom: ages > 90 days where fewer clones are represented removed.

| Response: Food consumed, 10^3^ cells / hour / individual | | | | |
| --- | --- | --- | --- | --- |
| Source | DF | Sum of Squares, x10^-3^ | F Ratio | Prob > F |
| Age | 1 | 96.56 | 12.56 | 0.0005 |
| Clone | 11 | 151.2 | 1.79 | 0.06 |
| Age*Clone | 11 | 142.82 | 1.69 | 0.08 |
| Error | 264 | 2030.35 |  |  |
| Response: Food consumed, 10^3^ cells / hour / mg WW | | | | |
| Source | DF | Sum of Squares, x10^-3^ | F Ratio | Prob > F |
| Age | 1 | 3.52 | 9.36 | 0.0024 |
| Clone | 11 | 20.05 | 4.85 | <0.0001 |
| Age*Clone | 11 | 5.28 | 1.28 | 0.24 |
| Error | 264 | 99.14 |  |  |

| Response: Food consumed, 10^3^ cells / hour / individual | | | | |
| --- | --- | --- | --- | --- |
| Source | DF | Sum of Squares, x10^3 | F Ratio | Prob > F |
| Age | 1 | 99.93 | 12.92 | 0.0004 |
| Clone | 11 | 147.21 | 1.73 | 0.07 |
| Age*Clone | 11 | 119.15 | 1.4 | 0.17 |
| Error | 254 | 1963.88 |  |  |
| Response: Food consumed, 10^3^ cells / hour / mg WW | | | | |
| Source | DF | Sum of Squares, x10^3 | F Ratio | Prob > F |
| Age | 1 | 3.61 | 9.47 | 0.0023 |
| Clone | 11 | 18.67 | 4.45 | <0.0001 |
| Age*Clone | 11 | 5.34 | 1.27 | 0.24 |
| Error | 254 | 96.93 |  |  |

A	B

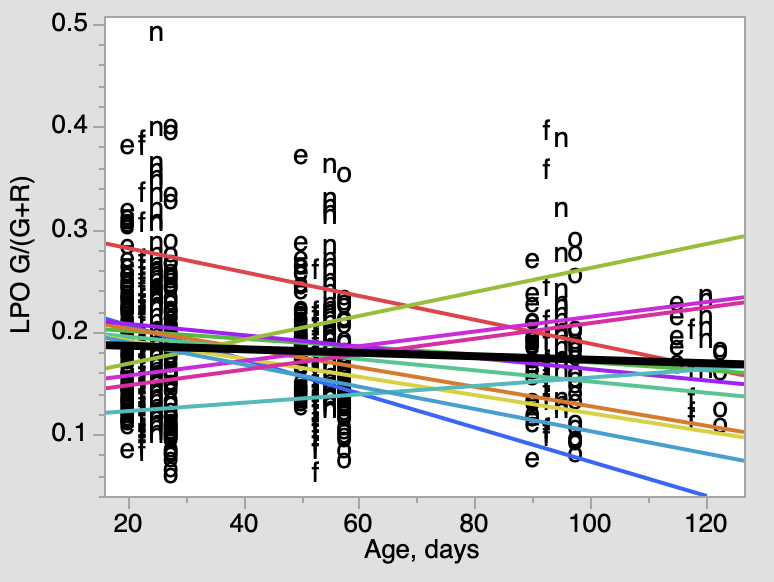

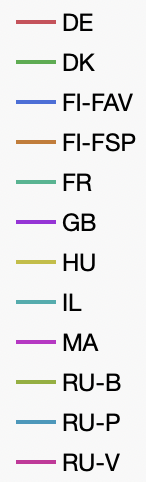

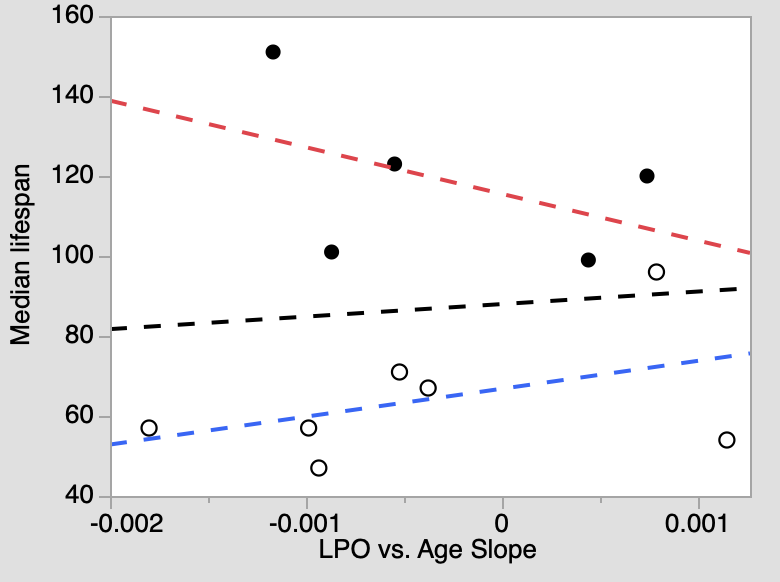

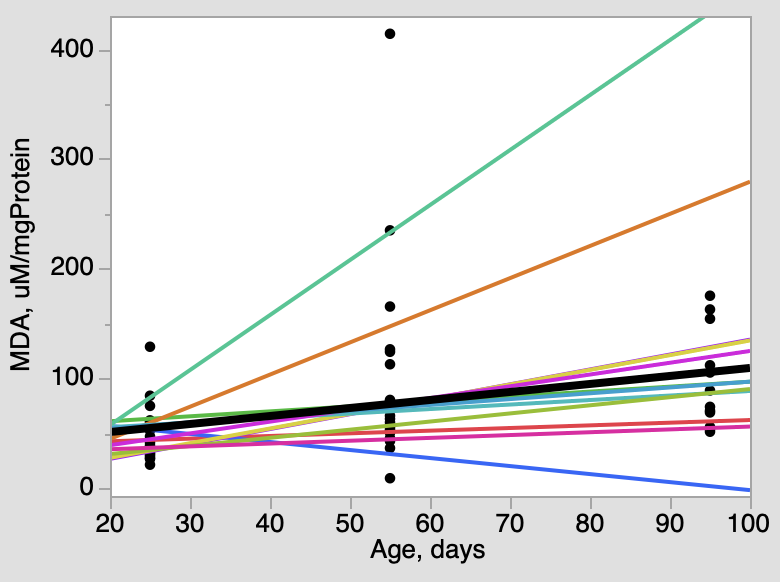

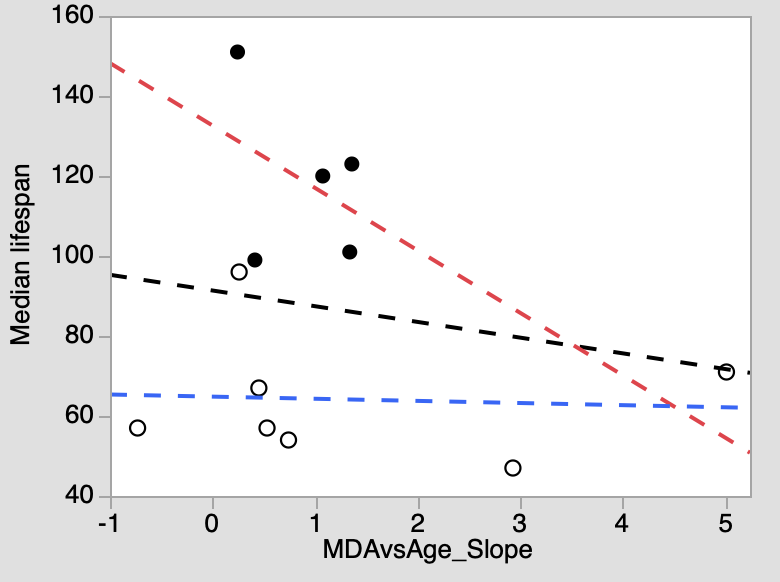

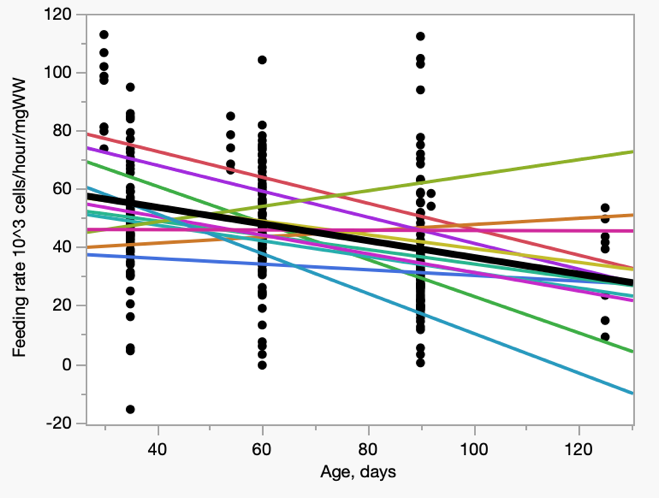

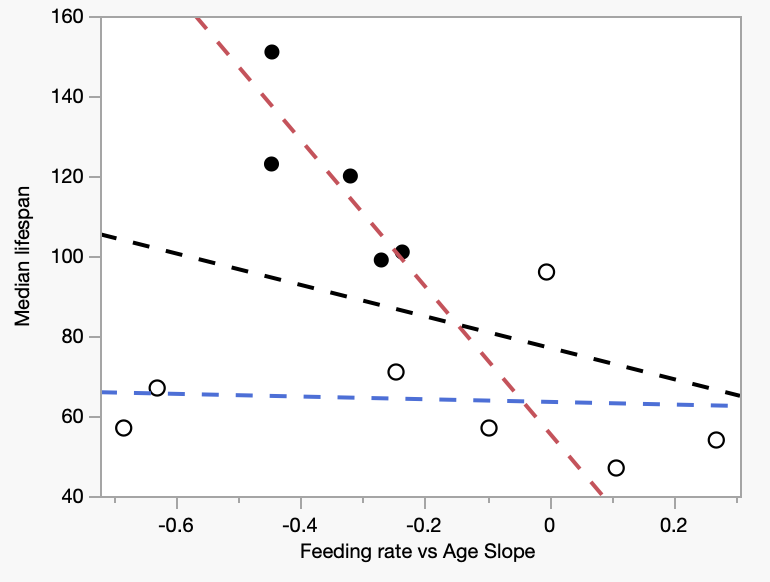

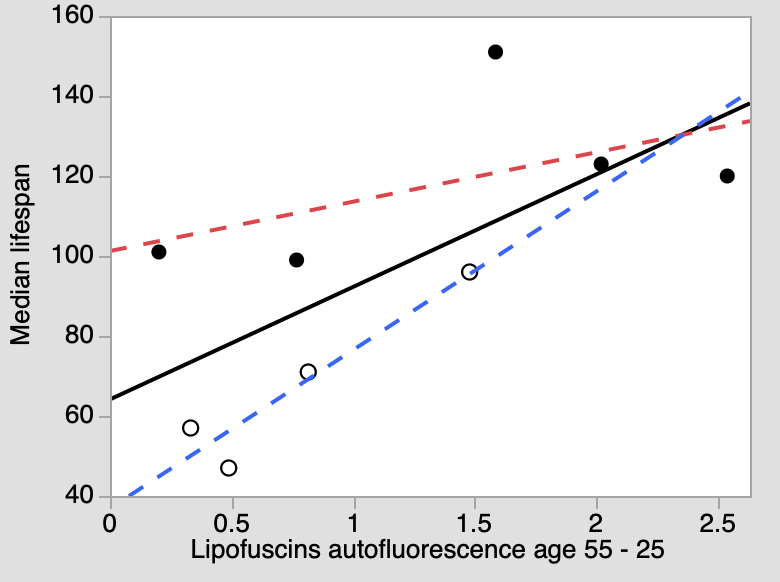

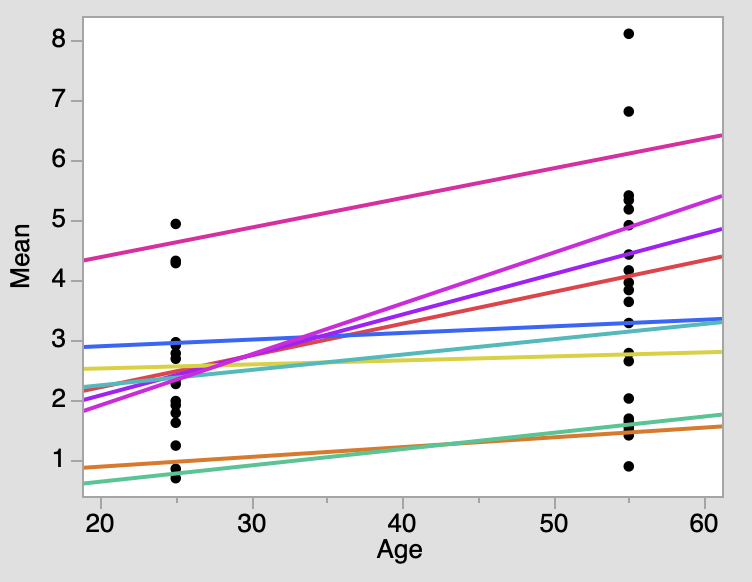

C	D

E	F

G	H

Fig. S6. Changes in feeding rate (10^3^ cells/hour, A), lipofuscins autofluorescence (C) and in two parameters of lipid peroxidation (LPO) with age in 12 clones of *D. magna* (E, G) and correlations between regression coefficients over age with clone-specific median lifespan (B, D, F, H)*.* Background-subtracted lipofuscins autofluorescence measured by fluorescence microscopy. LPO measured by fluorescence microscopy of *Daphnia* stained with BODIPY C11 dye, G/(G+R): ratio of median green fluorescence (G; oxidized dye) to the sum of G and median red fluorescence (R, reduced dye). Letters indicate body regions: e, epipodite-2; f, fat body; n, nephridium; o, ovary. Horizontal axis values staggered to show different letters. MDA (mM / mg protein) measured in whole body extractions. Regression lines on panels A, C, and E are drawn for all data (black lines) and separately for each clone (colored lines). Regression lines on panels B, D, and F are drawn for all 12 clones (black lines) and separately for the long-lived, caloric restriction sensitive and short-lived, caloric restriction insensitive clones (black and open symbols and red and blue lines, respectively). Regression lines on panels B, D, and F that are not statistically significant are shown dashed.

See Supplementary Tables S5, S6, and S7, respectively, for statistical analysis for panels A, C, and E.

Table S6. Effect of age (55 vs. 25 days) on while lipofuscins accumulation in 9 clones of *Daphnia.*

| Source | DF | Sum of Squares | F Ratio | Prob > F |
| --- | --- | --- | --- | --- |
| Clone | 8 | 50.96 | 9.42 | <0.0001 |
| Age | 1 | 16.45 | 24.31 | 0.0001 |
| Clone*Age | 8 | 2.88 | 0.53 | 0.82 |
| Error | 17 | 11.5 |  |  |

Table S7. ANOVA of the effects of age, clones, and body regions (ROI) on lipid peroxidation measured by fluorescence microscopy of *Daphnia* stained with BODIPY C11 dye. Age is treated as a continuous covariable. P-values <0.001 shown in bold. Top to bottom: entire data, absolute age; entire data, age normalized by clone-specific median lifespan; absolute age, only clones surviving to 90 days included; absolute age, all clones, but only age <90 included.

Absolute age, all clones, all ages

| Source | DF | Sum of Squares | F Ratio | Prob > F |
| --- | --- | --- | --- | --- |
| Clone | 11 | 0.626 | 22.06 | **<0.0001** |
| ROI | 3 | 0.111 | 14.32 | **<0.0001** |
| Clone*ROI | 33 | 0.041 | 0.48 | 0.99 |
| Age | 1 | 0.021 | 8.2 | 0.0043 |
| Clone*Age | 11 | 0.272 | 9.57 | **<0.0001** |
| ROI*Age | 3 | 0.005 | 0.61 | 0.61 |
| Clone*ROI*Age | 33 | 0.049 | 0.58 | 0.97 |
| Error | 640 | 1.651 |  |  |

Relative age, all clones, all ages

| Source | DF | Sum of Squares | F Ratio | Prob > F |
| --- | --- | --- | --- | --- |
| Clone | 11 | 0.315 | 11.11 | **<0.0001** |
| ROI | 3 | 0.097 | 12.56 | **<0.0001** |
| Clone*ROI | 33 | 0.054 | 0.63 | 0.95 |
| relAge | 1 | 0.031 | 11.94 | **0.0006** |
| Clone*relAge | 11 | 0.288 | 10.14 | **<0.0001** |
| ROI*relAge | 3 | 0.004 | 0.58 | 0.63 |
| Clone*ROI*relAge | 33 | 0.048 | 0.57 | 0.98 |
| Error | 640 | 1.651 |  |  |

Absolute age, clones surviving past 90 days, all ages

| Source | DF | Sum of Squares | F Ratio | Prob > F |
| --- | --- | --- | --- | --- |
| Clone | 7 | 0.575 | 31.51 | **<0.0001** |
| ROI | 3 | 0.221 | 28.26 | **<0.0001** |
| Clone*ROI | 21 | 0.023 | 0.42 | 0.99 |
| Age | 1 | 0.016 | 6.13 | 0.014 |
| Age*Clone | 7 | 0.241 | 13.19 | **<0.0001** |
| Age*ROI | 3 | 0.003 | 0.44 | 0.73 |
| Age*Clone*ROI | 21 | 0.042 | 0.77 | 0.75 |
| Error | 551 | 1.437 |  |  |

Absolute age, all clones, age classes under 90 days

| Source | DF | Sum of Squares | F Ratio | Prob > F | |
| --- | --- | --- | --- | --- | --- |
| Clone | 11 | 0.631 | 23.58 | | **<0.0001** |
| ROI | 3 | 0.118 | 16.14 | | **<0.0001** |
| Clone*ROI | 33 | 0.052 | 0.64 | | 0.94 |
| Age | 1 | 0.032 | 13.05 | | **0.0003** |
| Clone*Age | 11 | 0.281 | 10.5 | | **<0.0001** |
| ROI*Age | 3 | 0.018 | 2.45 | | 0.06 |
| Clone*ROI*Age | 33 | 0.03 | 0.38 | | 1 |
| Error | 519 | 1.262 |  | |  |

Table S8. ANOVA of the effects of age and clones on lipid peroxidation measured by fluorescence microscopy of *Daphnia* stained with BODIPY C11 dye in each body region separately. Age is treated as a continuous covariable. P-values <0.001 shown in bold. Cf. Supplementary Fig. S7.

| ROI: epipodite2 | |  |  |  |
| --- | --- | --- | --- | --- |
| Source | DF | Sum of Squares | F Ratio | Prob > F |
| Clone | 11 | 0.137 | 6.46 | **<0.0001** |
| Age | 1 | 0.001 | 0.36 | 0.55 |
| Clone*Age | 11 | 0.047 | 2.22 | 0.015 |
| Error | 163 | 0.313 |  |  |
| ROI: fat body |  |  |  |  |
| Source | DF | Sum of Squares | F Ratio | Prob > F |
| Clone | 11 | 0.158 | 5.84 | **<0.0001** |
| Age | 1 | 0.009 | 3.87 | 0.051 |
| Clone*Age | 11 | 0.078 | 2.91 | 0.0017 |
| Error | 153 | 0.375 |  |  |
| ROI: nephridium | |  |  |  |
| Source | DF | Sum of Squares | F Ratio | Prob > F |
| Clone | 11 | 0.197 | 5.48 | **<0.0001** |
| Age | 1 | 0.003 | 0.88 | 0.35 |
| Clone*Age | 11 | 0.084 | 2.35 | 0.0101 |
| Error | 166 | 0.541 |  |  |
| ROI: ovary |  |  |  |  |
| Source | DF | Sum of Squares | F Ratio | Prob > F |
| Clone | 11 | 0.173 | 5.89 | **<0.0001** |
| Age | 1 | 0.013 | 4.97 | 0.0271 |
| Clone*Age | 11 | 0.109 | 3.70 | **0.0001** |
| Error | 158 | 0.421 |  |  |

Table S9. ANOVA of the effects of age and clones on lipid peroxidation detox product MDA (mM/mg protein) measured in whole body extractions. Age is treated as a continuous covariable.

| Source | DF | Sum of Squares | F Ratio | Prob > F |
| --- | --- | --- | --- | --- |
| Clone | 11 | 73119.9 | 1.93 | 0.07 |
| Age | 1 | 24289.4 | 7.05 | 0.012 |
| Clone*Age | 11 | 29187.3 | 0.77 | 0.67 |
| Error | 36 | 123996.3 |  |  |

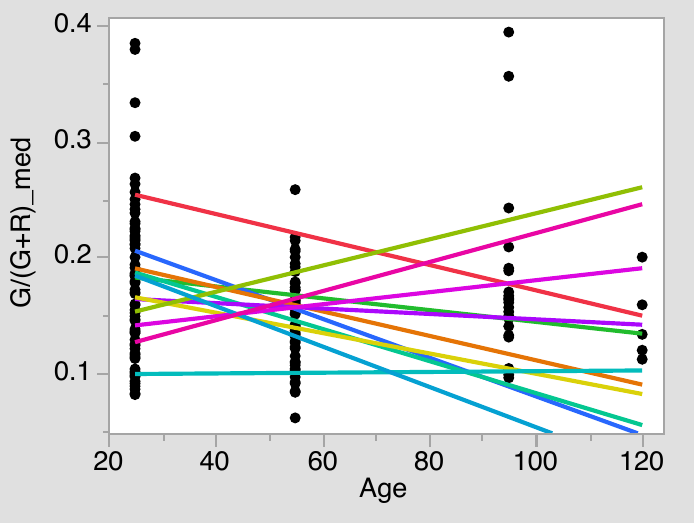

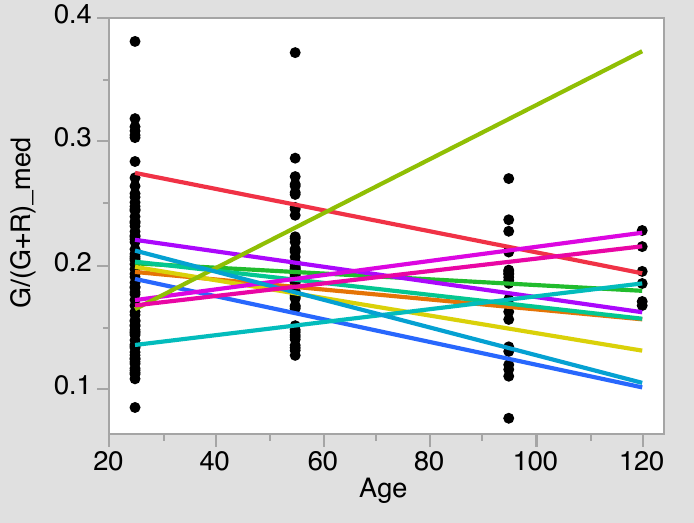
A	B

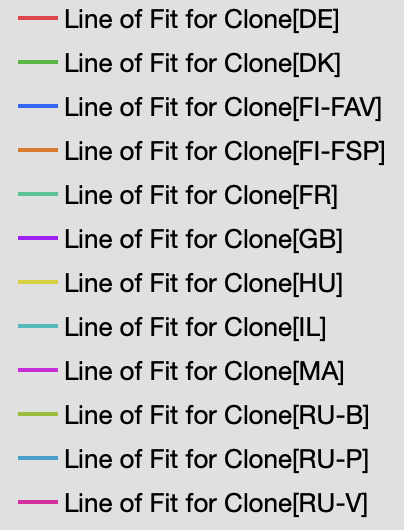

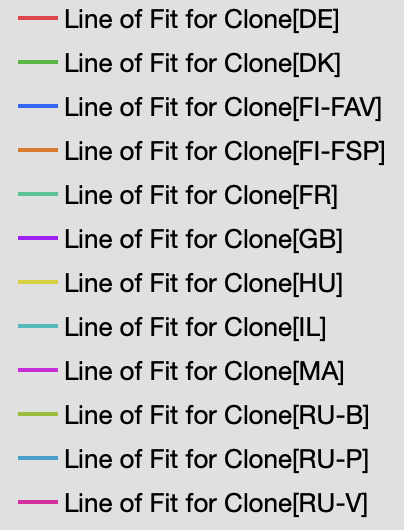

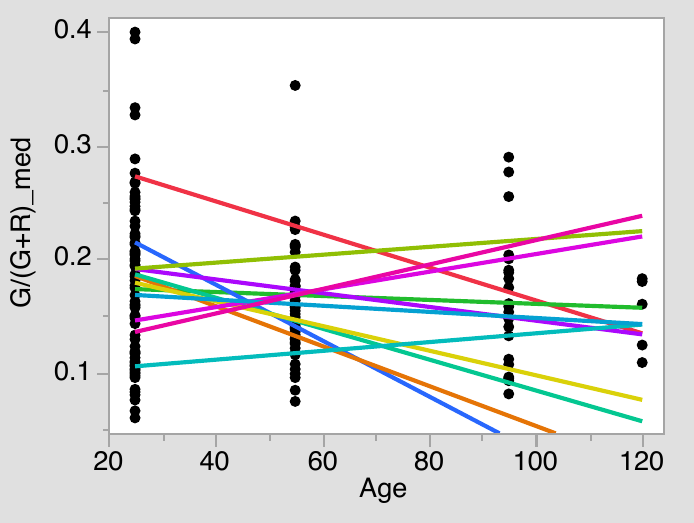

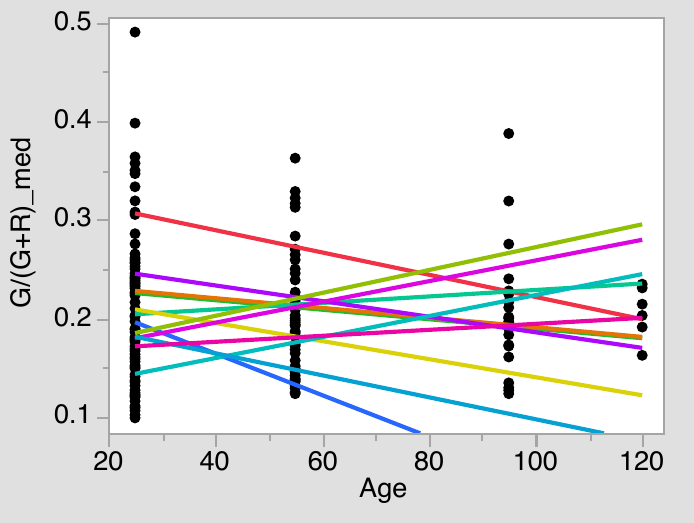

C	D

Supplementary Fig. S7. Changes of lipid peroxidation measured by fluorescence microscopy of *Daphnia* stained with BODIPY C11 dye in each body region. A: epipodite-2, B: fat body, C: nephridium, D: ovary. See Supplementary Table S’5’ for statistics.

Table S9. ANOVA of the effects of clonal means of feeding rate (top), LPO (middle), and MDA accumulation (bottom) measured in young a and middle-age Daphnia on clone-specific median lifespan. Main text Fig. 5-7.

| Source | DF | Sum of Squares | F Ratio | Prob > F |
| --- | --- | --- | --- | --- |
| Experiment | 1 | 8586.75 | 29.9 | <0.0001 |
| Clone Type | 1 | 2859.23 | 9.96 | 0.0031 |
| Feeding rate | 1 | 200.51 | 0.7 | 0.41 |
| Clone Type * Feeding rate | 1 | 44.63 | 0.16 | 0.70 |
| Age | 1 | 172.61 | 0.6 | 0.44 |
| Clone Type * Age | 1 | 71.11 | 0.25 | 0.62 |
| Feeding rate * Age | 1 | 118.99 | 0.41 | 0.52 |
| Clone * Feeding rate *Age | 1 | 230.1 | 0.8 | 0.38 |
| Error | 39 | 11200.93 |  |  |
| Source | DF | Sum of Squares | F Ratio | Prob > F |
| Experiment | 1 | 8586.75 | 40.1 | <0.0001 |
| Clone Type | 1 | 28828.6 | 134.62 | <0.0001 |
| LPO | 1 | 538.9 | 2.52 | 0.12 |
| Clone Type * LPO | 1 | 1507.11 | 7.04 | 0.012 |
| Age | 1 | 0.61 | 0.0029 | 0.96 |
| Clone Type * Age | 1 | 38.87 | 0.18 | 0.67 |
| LPO * Age | 1 | 185.97 | 0.87 | 0.36 |
| Clone Type * LPO *Age | 1 | 57.17 | 0.27 | 0.61 |
| Error | 39 | 8351.57 |  |  |
| Source | DF | Sum of Squares | F Ratio | Prob > F |
| Experiment | 1 | 8586.75 | 40.4 | <0.0001 |
| Clone Type | 1 | 8094.78 | 38.08 | <0.0001 |
| MDA | 1 | 579.79 | 2.73 | 0.11 |
| Clone Type * MDA | 1 | 454.51 | 2.14 | 0.15 |
| Age | 1 | 947.54 | 4.46 | 0.042 |
| Clone Type * Age | 1 | 850.27 | 4 | 0.053 |
| MDA * Age | 1 | 1996.31 | 9.39 | 0.004 |
| Clone Type * MDA *Age | 1 | 1910.55 | 8.99 | 0.005 |
| Error | 39 | 8289.95 |  |  |

Age = 25 days	Age = 55 days

epipodite2

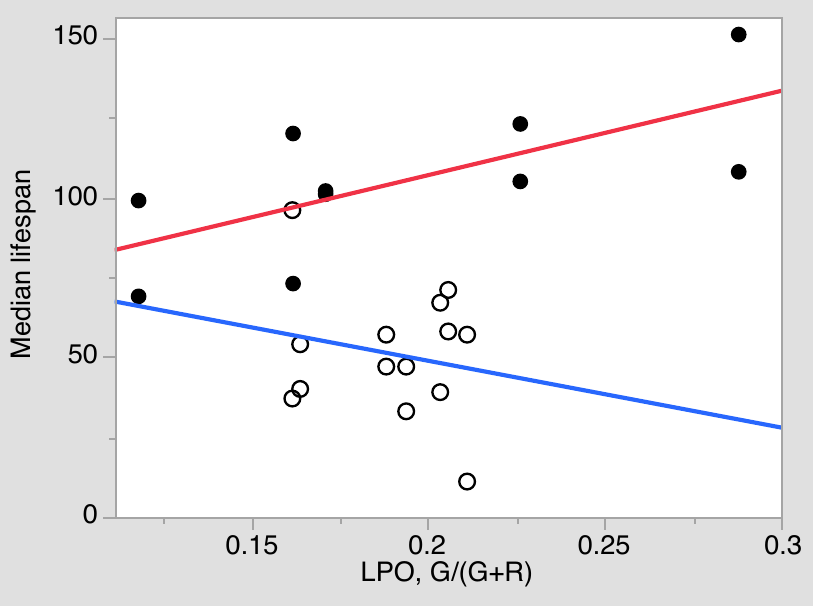

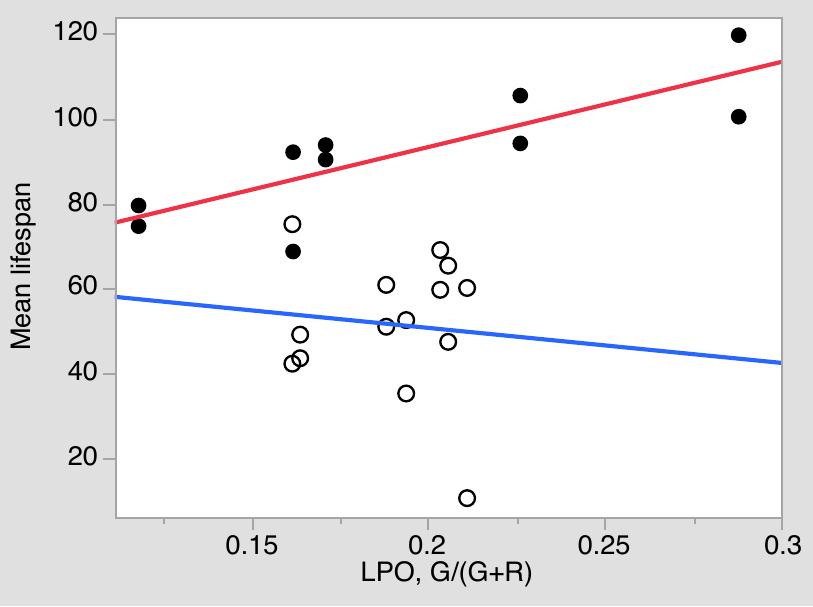

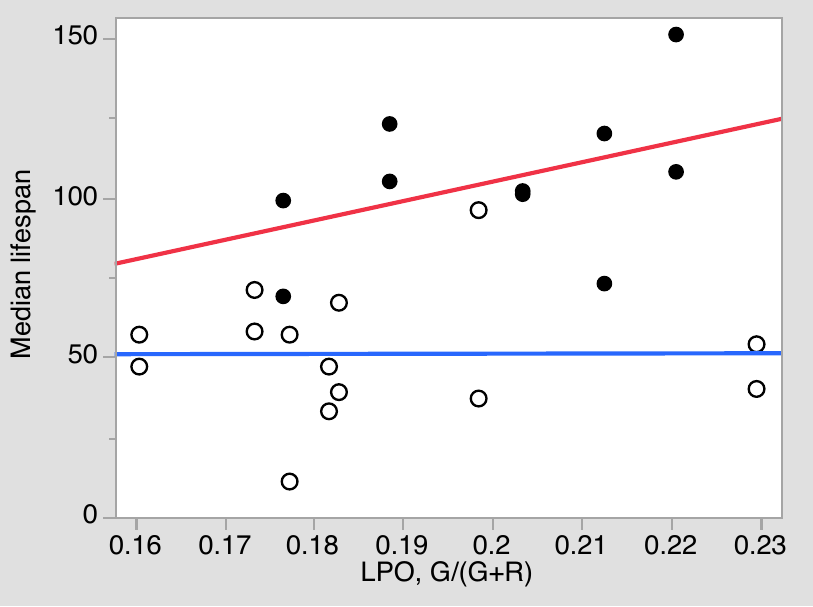

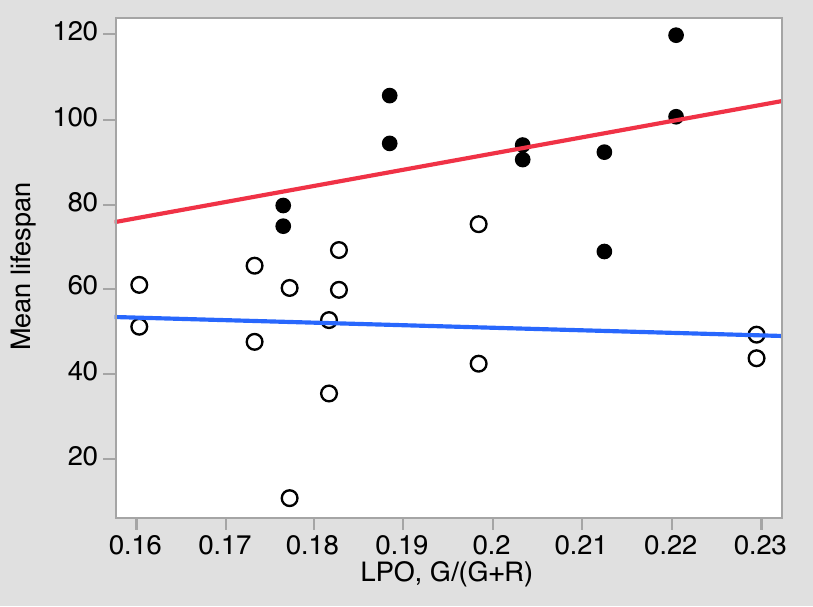

fat body

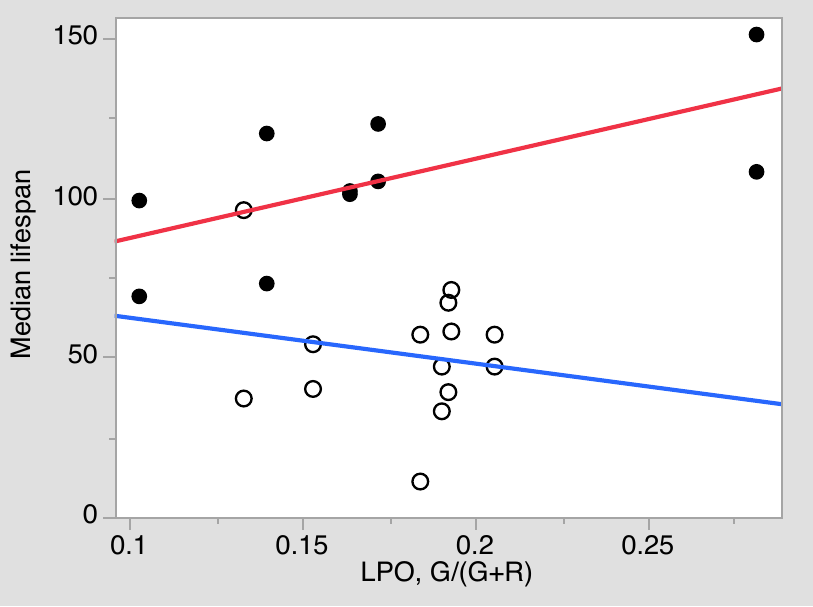

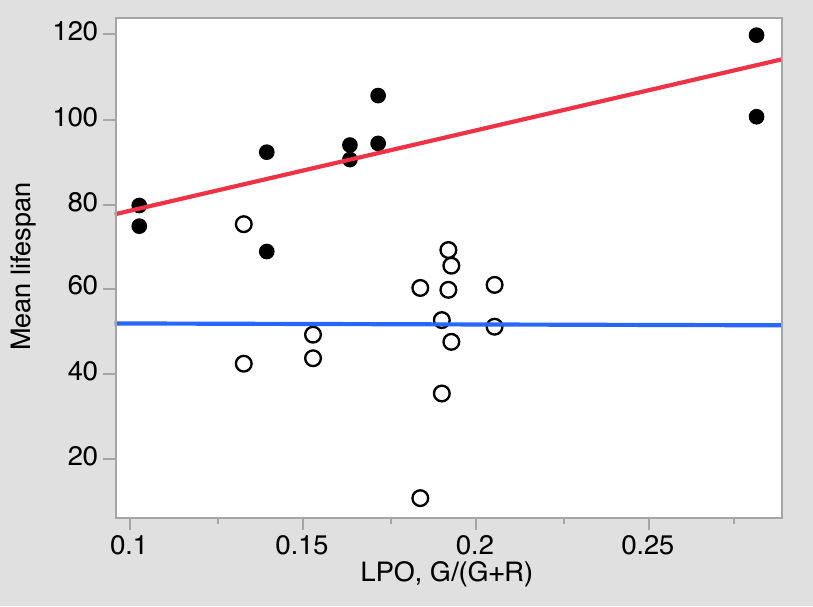

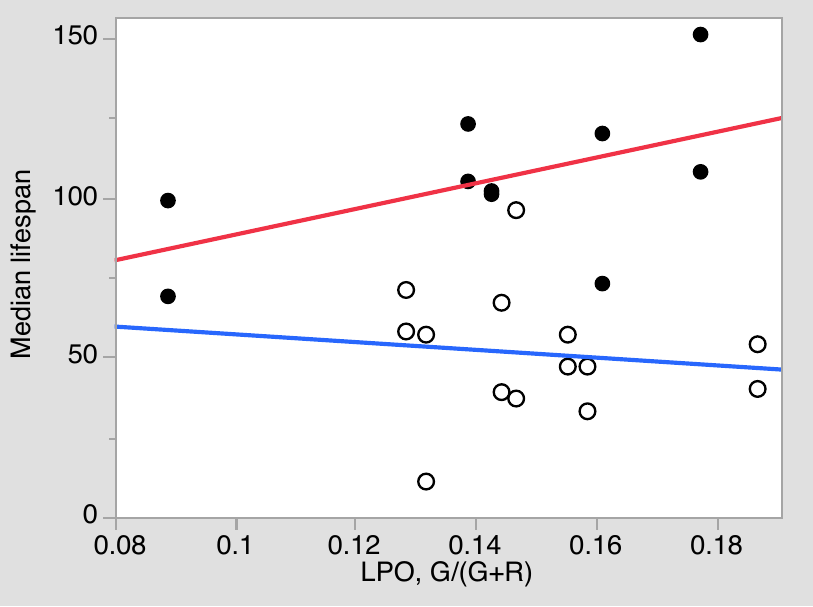

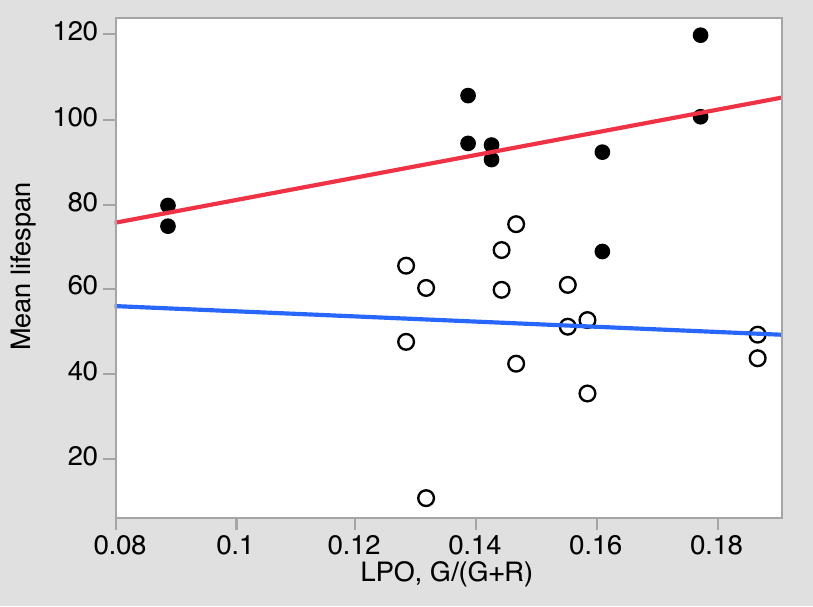

nephridium

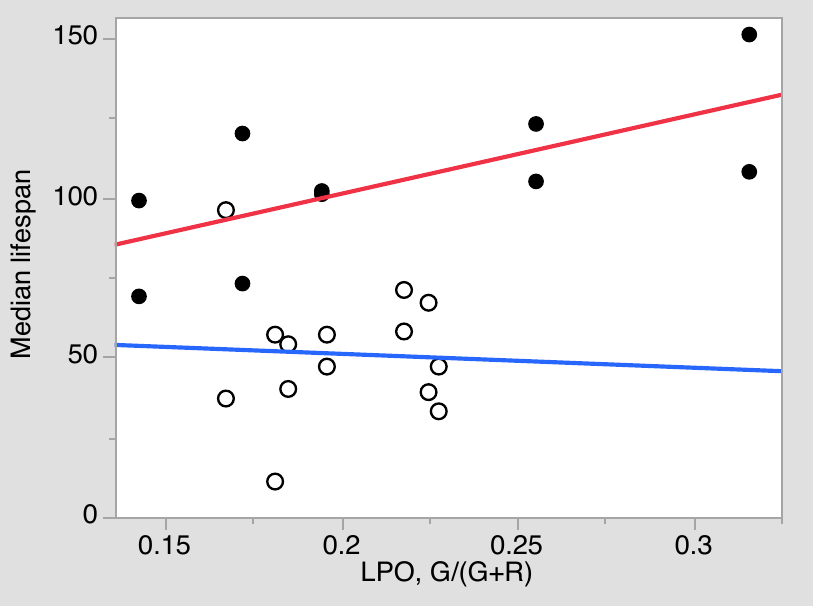

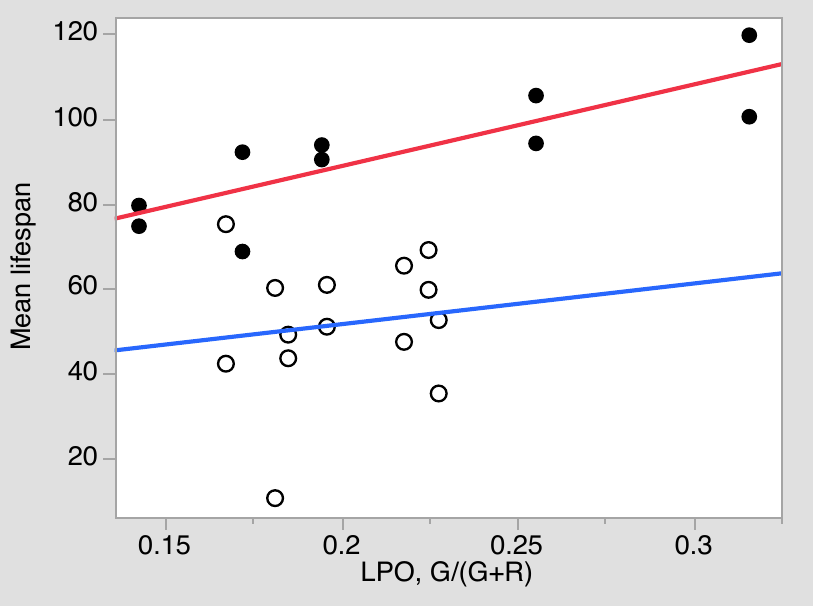

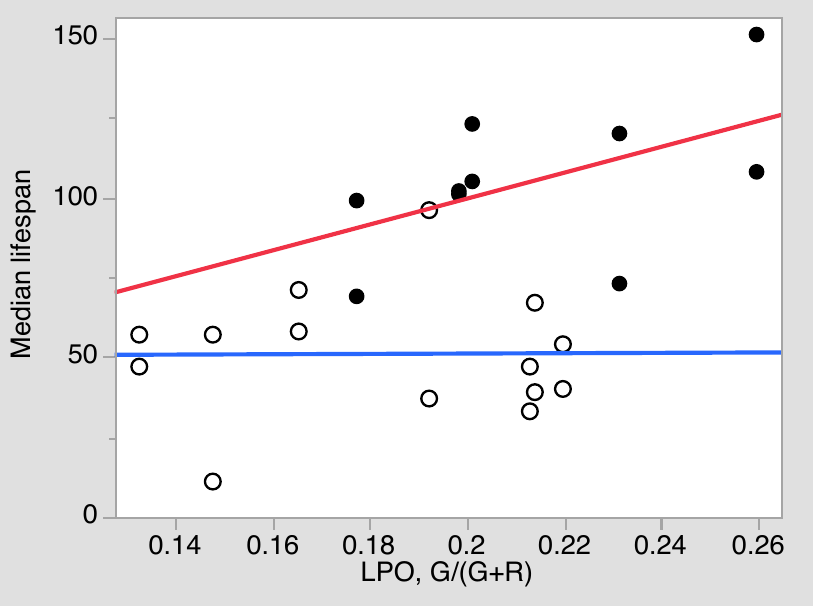

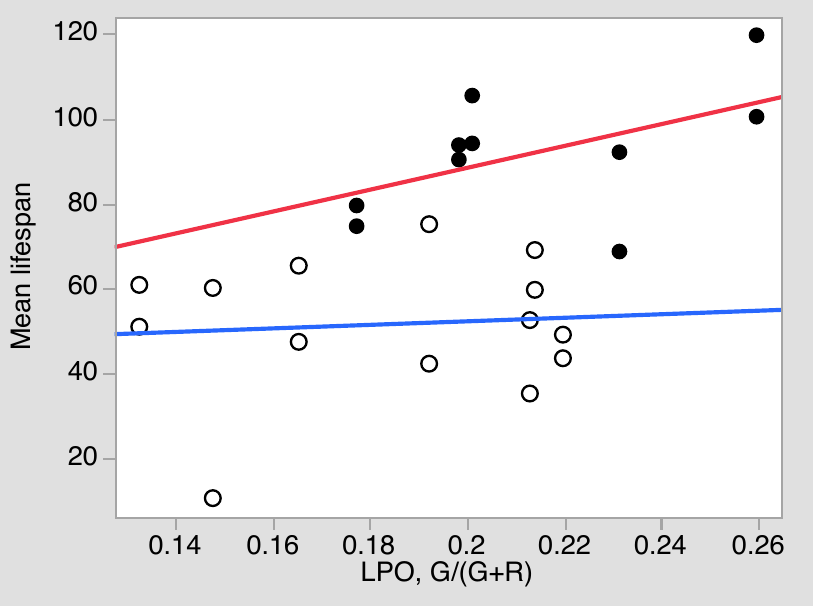

ovary

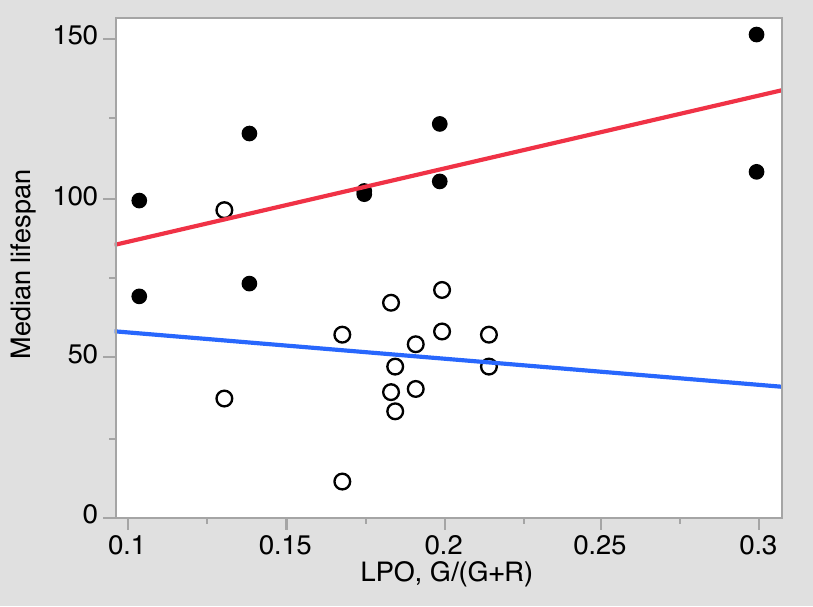

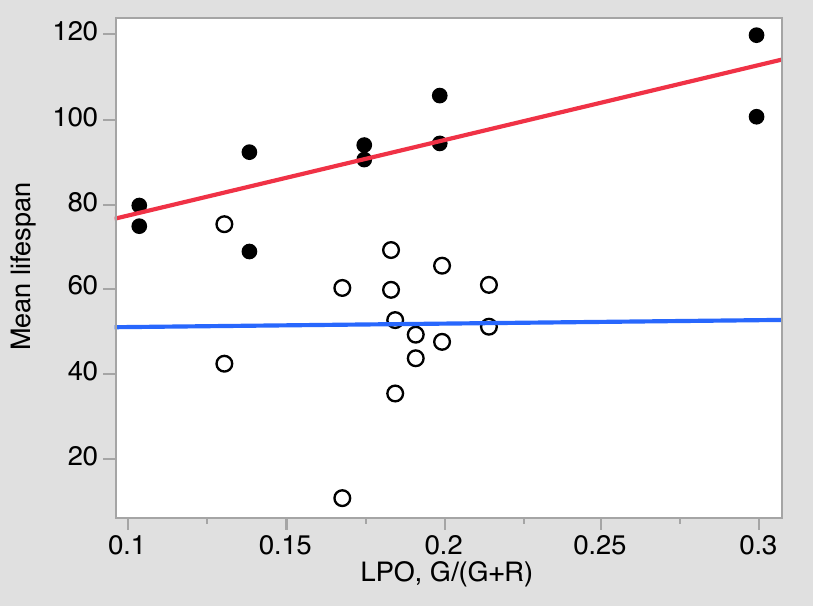

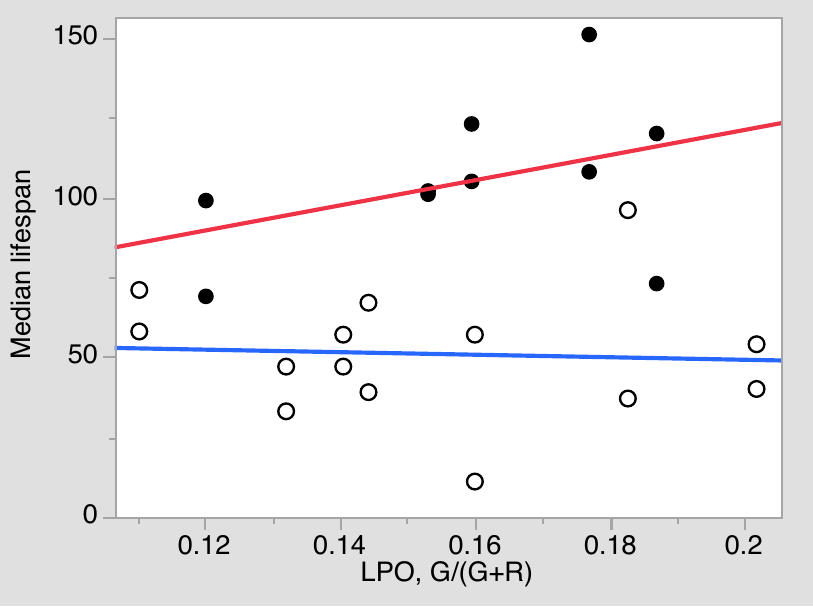

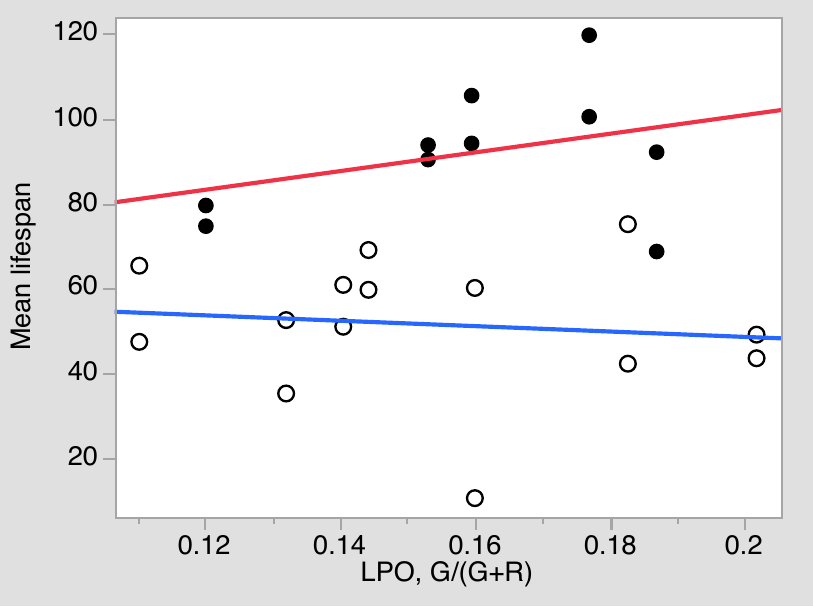

Fig. S8. As in main text Fig. 5 C, D, but each body part (ROI) considered separately.
